# The Janus-like role of neuraminidase isoenzymes in inflammation

**DOI:** 10.1101/2021.07.14.452400

**Authors:** Md. Amran Howlader, Ekaterina P. Demina, Suzanne Samarani, Tianlin Guo, Ali Ahmad, Alexey V. Pshezhetsky, Christopher W. Cairo

## Abstract

The processes of activation, extravasation, and migration of immune cells to a site are early and essential steps in the induction of an acute inflammatory response. These events are part of the inflammatory cascade which involves multiple regulatory steps. Using a murine air-pouch model of inflammation with LPS as an inflammation inducer we demonstrate that isoenzymes of the neuraminidase family (NEU1, 3, and 4) play essential roles in this process acting as positive or negative regulators of leukocyte infiltration. Genetically knocked-out (KO) mice for different NEU genes (*Neu1* KO, *Neu3* KO, *Neu4* KO, and *Neu3/4* double KO mice) were induced with LPS, leukocytes at the site of inflammation were counted, and the inflamed tissue was analyzed using immunohistochemistry. Our data show that leukocyte recruitment was decreased in NEU1 and NEU3-deficient mice, while it was increased in NEU4-deficient animals. Consistent with these results, systemic levels of pro-inflammatory cytokines and those in pouch exudate were reduced in *Neu1* and increased in *Neu4* KO mice. We found that pharmacological inhibitors specific for NEU1, NEU3, and NEU4 isoforms also affected leukocyte recruitment. We conclude that NEU isoenzymes have distinct – and even opposing – effects on leukocyte recruitment, and therefore warrant further investigation to determine their mechanisms and importance as regulators of the inflammatory cascade.

## 1 Introduction

The induction of the inflammatory cascade by exogenous or endogenous stimuli culminates in acute inflammation: a host protective response that tends to neutralize the stimulus and maintains tissue homeostasis and integrity. However, when acute inflammation becomes uncontrolled and long-lasting, it contributes to the pathogenesis of chronic inflammatory diseases such as sepsis, diabetes, atherosclerosis, and cardiovascular disease. The cascade begins with activation and trafficking of leukocytes to the site of inflammation.(Ley *et al*, 2007) This process provides a rapid response to an infection by regulating permeability of the vascular compartment, activation of endothelial cells, tethering and adhesion of leukocytes to the vascular endothelium, and subsequent extravasation to the site of inflammation as well as activation of macrophages, platelets, complement, and clotting factors. Acute inflammation can be triggered by recognition of pathogen-associated molecular patterns (PAMPs), such as the lipopolysaccharide (LPS) of gram negative bacteria, by one or more pattern recognition receptors (PRR) of the host cells (e.g. Toll-like receptors, TLRs).(Rossol *et al*, 2011) For example, LPS is recognized by TLR-4 leading to initiation of the inflammatory cascade and leukocyte recruitment.(Miyake, 2004) The glycosylation state and activity of cellular receptors, including TLR-4, is known to be regulated by endogenous neuraminidase enzymes (NEU, or sialidases).(Amith *et al*, 2010; Feng *et al*, 2012; Pshezhetsky & Hinek, 2011) Notably, there are four NEU isoenzymes, each with distinct tissue expression, sub-cellular localization, and roles in inflammation and pathogenesis.(Miyagi & Yamaguchi, 2012) These enzymes remove sialic acid from the termini of glycoproteins, glycolipids, and oligosaccharides in a substrate- and linkage-specific manner. Increased endogenous neuraminidase activity, accompanied by leukocyte activation, is known to play a role in the infiltration of leukocytes to the site of inflammation; however, the precise role of individual NEU isoenzymes in leukocyte recruitment has not been investigated in a systematic fashion.(Feng *et al*, 2011; Gadhoum & Sackstein, 2008)

*N*-Acetyl-neuraminic acid (Neu5Ac, or sialic acid) residues are known to be important in multiple steps of the inflammatory cascade. Selectins, expressed by immune cells as well as by vascular endothelial cells (VEC), mediate leukocyte capture and rolling and require minimal tetrasaccharide epitopes containing Neu5Ac residues (sialyl-Lewis^X^; sLe^X^ or CD15s) for binding with their ligands (e.g. PSGL-1, CD44).(Zen *et al*, 2007) Endogenous NEU are able to desialylate these epitopes, thus disrupting selectin-ligand interactions.(Gadhoum & Sackstein, 2008) The firm adhesion step of the cascade is mediated by activated leukocyte β2 integrins, LFA-1 (CD11a/CD18) and MAC-1 (CD11b/CD18), which bind immunoglobulin-like adhesion molecules (ICAM-1 and ICAM-2) expressed by activated VEC. Activation epitopes of β2 integrins were reported to be unmasked by endogenous NEU activity,(Feng *et al.*, 2011; Quinn *et al*, 2001) and NEU3 has been shown to alter LFA-1/ICAM-1 interactions.(Howlader *et al*, 2019) NEU enzymes can increase adhesion of polymorphonuclear leukocytes (PMN) to endothelium and also increase their migration through the endothelium to the site of inflammation.(Cross *et al*, 2003; Sakarya *et al*, 2004) NEU enzymes may unmask epitopes in β1 integrins VLA4 (CD49d/CD29) and VLA5 (CD49e/CD29), which are expressed on activated leukocytes and bind with VCAM-1 and fibronectin (FN).(Feng *et al.*, 2011; Quinn *et al.*, 2001) Activated PBMCs have been shown to increase production of inflammatory cytokines such as TNF-α and IFN-γ through the action of NEU1 or NEU3.(Nan *et al*, 2007) The effect of cytokine production (e.g. IL-6, IL-12p40, and TNF-α) in dendritic cells has been reported to be regulated by NEU1 and NEU3 activity.(Stamatos *et al*, 2010) NEU1 has been shown to induce phagocytosis in macrophages by activation of Fc-γ receptors.(Seyrantepe *et al*, 2010) Together, these findings suggest that endogenous NEU activity is involved at multiple points along the inflammatory cascade.

To test the hypothesis that individual NEU isoenzymes have different effects on inflammation and to address specific roles of these enzymes in recruitment of leukocytes to the site of inflammation, we used a murine six-day air pouch model of LPS-induced acute inflammation(Sin *et al*, 1986) in *Neu1*, *Neu3*, and *Neu4* KO and WT mice (in an identical C57Bl6 genetic background). In previous studies we demonstrated that *Neu3* and *Neu4* enzymes have similar substrate specificity and can complement each other;(Smutova *et al*, 2014) therefore, we also tested *Neu3/4* double KO (DKO) animals deficient in both enzymes.(Pan *et al*, 2017) We observed that NEU isoenzymes drastically modulated the inflammatory response to LPS, with NEU1 and NEU3 isoenzymes acting as positive regulators and NEU4 acting as a negative regulator of the response. These results reveal that NEU isoenzymes have important and distinct roles in the regulation of immune cell migration to the site of inflammation as well as consequent inflammation. Moreover, our findings suggest that NEU isoenzymes play roles in immune response and may be targets to modulate pathogenic inflammation in human disease.

## 2 Materials and Methods

### 2.1 Animal models

C57BL6 mice (aged 3-4 months) were used as wild type control. *Neu1* KO, *Neu3* KO, *Neu4* KO, and *Neu3/4* DKO mice were as reported previously.(Pan *et al.*, 2017) Mice were housed in an enriched environment with continuous access to food and water, under constant temperature and humidity, on a 12 h light/dark cycle. The experiments involving animals were approved by the Animal Care and Use Committee of the CHU Ste-Justine Research Center (protocol #710). WT and *Neu1* KO group contained both male and female mice, while other groups contained male mice only.

### 2.2 Sources of reagents and antibodies

Compounds **IN1**(CG14600), **IN3**(CG22600), and **IN4**(CY16600) were prepared as previously reported.(Albohy *et al*, 2013; Guo *et al*, 2018a; Guo *et al*, 2018b; Zhang *et al*, 2013) Stock solutions for all compounds were made using saline. An LPS isolated from *E. coli* O55:B5 that does not contain sialic acid was purchased from Sigma Aldrich (cat# L2880).(Lindberg *et al*, 1981) Anti-mouse/human CD11b FITC (Clone M1/70), anti-mouse CD45 Alexa Fluor 700 (Clone 30-F11), anti-mouse/human CD45R/B220 PerCP (Clone RA3-6B2), anti-mouse 49b PE (Clone DX5), anti-mouse F4/80 APC (Clone BM8), anti-mouse Ly-6G PE/Cyanine 7 (Clone 1A8), anti-mouse Ly-6C Brilliant violet 421 (Clone HK1.4), and anti-mouse CD115 APC/Cyanine 7 (Clone AFS98) were purchased from Biolegends, USA.

### 2.3 Air pouch model of acute inflammation

The air pouch inflammation model was previously described and used here with slight modifications.(Cronstein *et al*, 1993; Terkeltaub *et al*, 1998; Tessier *et al*, 1997) Briefly, mice (6-8 weeks, male or female) were randomly separated into control and experimental groups. Hair was removed at the dorsal area (5×2 cm) two days prior to commencing the experiment, vaseline was applied on the nude area to alleviate discomfort. Animals were monitored for abnormal behavior and skin injuries daily. On days 3 and 6, mice were anesthetized under isoflurane and 3 mL of sterilized air (passed through a 0.2 μm filter) was injected subcutaneously into the back of the mice using a 26-gauge needle. For treated mice, an intraperitoneal injection of an inhibitor on days 6, 7, and 8 (200 μL of saline containing inhibitors at 1 mg/kg). Saline was injected for control groups and mice that were not subjected to inhibitors. On day 8, 1 mL of sterile PBS or sterile PBS with LPS (1 μg/mL) was injected into the air pouches. At 9 h post-injection, mice were sacrificed using phenobarbital overdose (150 mg/kg BW). Then air pouches were washed with HBSS containing 10 mM EDTA (1 mL, 2 × 2 mL). The exudates were collected and centrifuged at 100 × g for 10 min at room temperature. The supernatants were collected and frozen for later analysis. Cells were re-suspended in 1 mL of HBSS-EDTA and counted by haemocytometer or used in flow cytometry analysis.

### 2.4 Flow cytometry analysis of immune cells

Subpopulations of cells isolated from the air pouch exudate were analyzed by flow cytometry as previously described.(Nguyen *et al*, 2013) Dead cells were stained with Aqua blue (Thermo Fisher Scientific) and Fc Receptors (FcR) were blocked using mouse IgG. After washing with PBS (containing 2% FBS), cells were incubated on ice for 30 min with fluorophore-conjugated antibodies as a mixture: CD49b PE, CD45 AF700, CD115 APC/CY7, Ly-6C BV421, Ly-6G PE/CY7, F4/80 APC, B220 PerCP, CD11b FITC, CD3 FITC, and CD11c APC. Briefly, cells were washed twice with PBS, resuspended in 2% paraformaldehyde, and analyzed by Flow cytometry using BD LSRII Fortessa. Single fluorochromes were used for compensation (elimination of spectral overlap), and for setting of gates using a minus one method. Data were analyzed by Diva and FlowJo. Dead cells and doublets were eliminated from the analyses.

### 2.5 Immunohistochemistry protocol and cell counts

Immunohistochemistry (IHC) slides of Tissue Tek OCT pouch tissue were prepared by making 4 μm thick slices and then staining them with hematoxylin and eosin (H&E) stain. The slides were then imaged using a ZEISS Axio Scan Z1. Regions of the IHC slides were identified as three major regions: epidermis layer, dermis, and underlying muscle layer. The H&E stain revealed leukocytes present in each region. The number of leukocytes were counted with ImageJ using at least three randomly chosen fields of the same region. Skin sections of at least three mice were investigated from each group.

### 2.6 Cytokine analysis using ProcartaPlex immunoassays

Levels of cytokines in mouse plasma and exudates were investigated using ProcartaPlex immunoassays (Perkin Elmer, USA) with a custom panel of 21 cytokines according to the manufacturer’s protocol. Briefly, plasma and exudate samples were diluted in sample diluent and incubated with capture antibody-coupled magnetic beads on a shaker at 4 °C overnight. After removal of samples by centrifugation and three washes, biotinylated secondary antibody was added for incubation on shaker in the dark at room temperature for 30 min. Each captured cytokine was detected by addition of streptavidin-phycoerythrin and fluorescence was measured using the Autoplex Analyser CS1000 system (Perkin Elmer, Waltham, MA).

### 2.7 Macrophage preparation and chemotaxis trans-migration studies

Monocytes (MO) were isolated from the bone marrow of mice and differentiated in vitro into macrophages (MΦ) by incubating bone marrow cells in complete DMEM (cDMEM, DMEM supplemented with 10% heat-inactivated FBS, 10% conditioned media supernatant from L-929 fibroblast cell cultures, 1% penicillin-streptomycin, 0.01M HEPES buffer, 1 mM sodium pyruvate). Macrophages were cultured for 7 days in a T75 flask with 80% confluency. Cells were then used for a transmigration assay adopted from a previous study.(Senger *et al*, 2002) Briefly, an 8-μm-pore size transwell migration plate (Costar, Fischer Sci, USA) was used after the transwell membrane was coated with 50 μg/mL fibronectin (FN) in PBS for 1 h and blocked with bovine serum albumin (BSA) for 30 min. The bottom chamber of the plate contained PBS with 20 ng/well monocyte chemoattractant protein-1 (MCP-1, Thermofisher Scientific, USA). Macrophages (5×10^4^ cells in PBS) isolated from the genotypic mice were carefully placed on the top chamber and incubated for 5 h at 37 °C with 5% CO_2_. Subsequently, the bottom of the upper chamber was carefully wiped using a Q-tip and the transwell membrane was stained. Cells that had infiltrated the membrane were then imaged using Zeiss COLIBRI fast LED imaging and optical sectioning microscope using 10X lens under bright field. The images were analyzed using Zen black software (Zeiss).

## 3 Results

### 3.1 Inflammatory response to LPS reveals involvement of neuraminidase enzymes in the recruitment of leukocytes

We used a murine air-pouch model to investigate the effect of individual neuraminidase enzymes on LPS-induced inflammation.(Sin *et al.*, 1986) An air pouch was created by subcutaneous injection of sterile air into the lower dorsal region of WT and *Neu1* KO, *Neu3* KO, *Neu4* KO, and *Neu3/4* DKO mice (see Materials & Methods). The pouch was injected with either 1 μg LPS in 1 mL of saline, to simulate a bacterial infection, or 1 mL of saline, which served as a control for basal levels of inflammation. After an incubation period of 9 h, the air pouch was washed with sterile saline to harvest cells, which were then counted by flow cytometry (**Figure 1, Table 1**). The differences in the pouch cell counts between WT and the *Neu* KO mice at basal level were non-significant (**Figure 1B**). In contrast, LPS treatment showed significant differences in the cell numbers between WT and *Neu* KO mice (**Figure 1C**). In the case of *Neu1* KO animals, leukocyte counts were reduced to ~25% (79 ± 9 × 10^4^ cells/pouch) of those in WT mice (326 ± 29 × 10^4^ cells/pouch). The *Neu3* KO animals also showed a trend for reduction of leukocytes infiltrating the pouch in response to LPS as compared to the WT mice; however, the difference was not statistically different from control. The *Neu4* KO (1371 ± 229 × 10^4^ cells/pouch) and *Neu3/4* DKO (992 ± 141 × 10^4^ cells/pouch) animals demonstrated significant increases (3-4 fold) in cell numbers as compared with the WT mice upon LPS treatment. These results suggested that LPS-stimulated leukocyte infiltration into the pouch was positively regulated by NEU1, and negatively regulated by NEU4.

**Figure 1.**
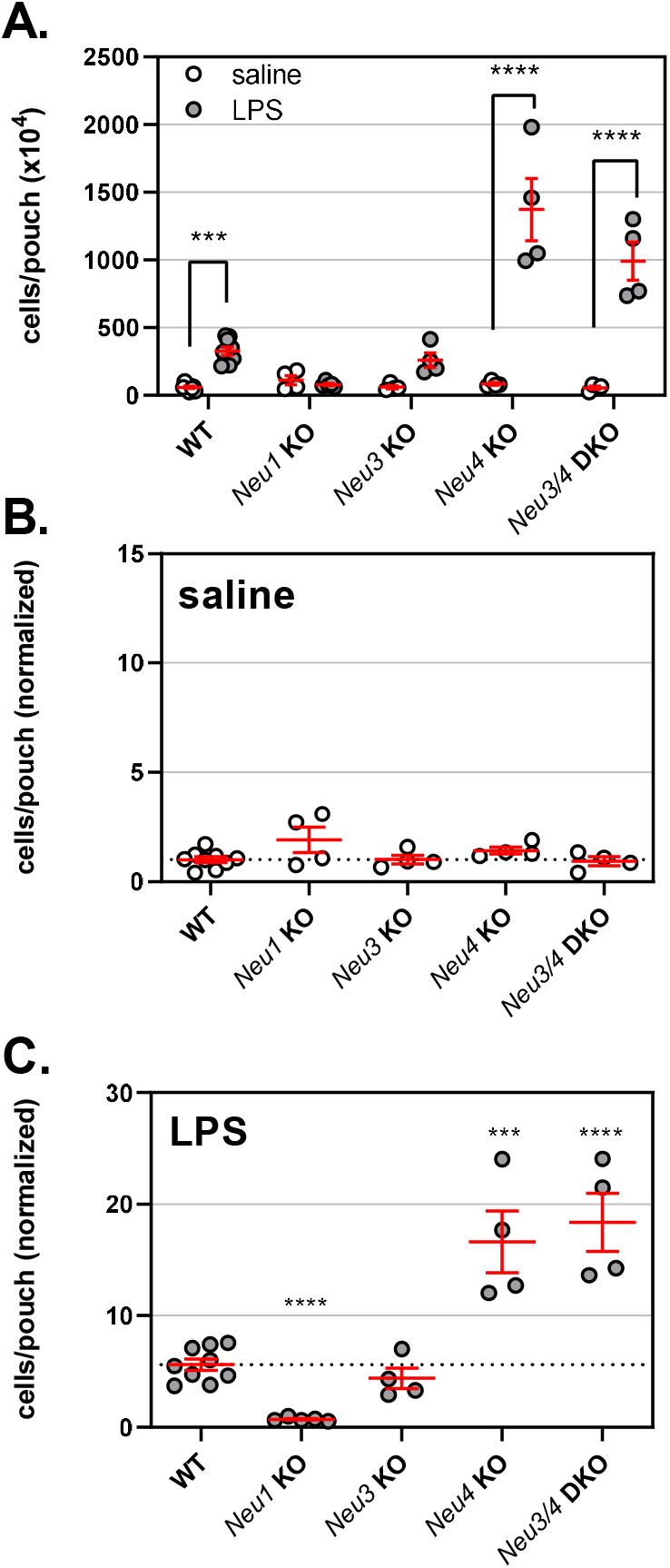
Leukocyte counts from the air pouch exudate. An air pouch was formed after injection of sterile air for each genotype. The pouch was then injected with saline or LPS and incubated for 9 h. Mice were then sacrificed, the pouch exudate was collected, and cells were counted by flow cytometry. A. Cell counts are presented for the saline (○) and LPS (●) treatments. B. Cell counts for the saline treatment in the *Neu* KO mice are compared with those of WT mice (58 × 10^4^ cells/pouch; normalized to 1, dashed line). C. Cell counts for the LPS treatment are compared with those of WT mice (normalized to WT saline treatment). Individual points are shown with mean ± SEM. For panel A, comparisons were made using two-way ANOVA following Dunnet’s multiple comparison test; for panel B & C, comparisons were made using one-way ANOVA and Dunnet’s t-test to compare groups to control. (*, p ≤ 0.05; **, p ≤ 0.01; ***, p ≤ 0.005; ****, p ≤ 0.0001).

**Table 1.** Leukocyte raw cell counts observed in the air pouch model.

|  | Saline<br>(cells x10<sup>4</sup>) |  |  | LPS<br>(cells x10<sup>4</sup>) |  |  |
| --- | --- | --- | --- | --- | --- | --- |
|  | Mean ± SEM | *p* | N | Mean ± SEM | *p* | N |
| **WT C57BL6** | 58 ± 8 |  | 9 | 326 ± 29 |  | 9 |
| ***Neu1* KO** | 111 ± 34 |  | 4 | 79 ± 9 | **** | 5 |
| ***Neu3* KO** | 59 ± 12 |  | 4 | 259 ± 54 |  | 4 |
| ***Neu4* KO** | 83 ± 9 |  | 4 | 1371 ± 229 | *** | 4 |
| ***Neu3/4* DKO** | 54 ± 12 |  | 4 | 992 ± 141 | **** | 4 |
| **WT IN1** | 72 ± 16 |  | 4 | 357 ± 111 |  | 4 |
| **WT IN3** | 100 ± 26 |  | 4 | 189 ± 40 |  | 4 |
| **WT IN4** | 172 ± 43 | ** | 4 | 433 ± 49 |  | 3 |

Values were compared to control using a Student's t-test (\*, $p \leq 0.05$ ; \*\*, $p \leq 0.01$ ; \*\*\*, $p \leq 0.005$ ; \*\*\*\*, $p \leq 0.0001$ ).

### 3.2 Leukocyte subset response to LPS in neuraminidase deficient animal models

To determine the influence of neuraminidases on leukocyte subsets found in the air pouch model, we stained cells from the exudate with cell type-specific antibodies and quantified them using flow cytometry. Cell counts were normalized to saline-treated WT control for the selected leukocyte subsets shown in **Figure 2** (**Figure SI1** and **Table SI1** provide raw counts). We observed that the major populations found in the air pouch after LPS treatment were monocytes (MO), neutrophils (NE), natural killer cells (NK), and macrophages (MΦ). T and B lymphocyte counts were low for both conditions, and no significant changes in their levels were detected for any of NEU-deficient animals as compared to WT mice (**Table SI1**). Other subsets analyzed revealed cell type-specific differences between NEU-deficient and WT mice. Specifically, *Neu1* KO animals had increased basal MO and NE counts (saline treatment). In contrast, *Neu4* KO animals had elevated counts of MO, NE, and NK cells after LPS treatment (3-9 fold increase as compared to WT), as well as elevated basal MO counts (5-fold increase as compared to WT). The *Neu3* KO animals had elevated MO counts in saline controls. The *Neu3/4* DKO animals showed moderate elevation of MO counts after LPS treatment. These results are generally consistent with NEU4 acting as a negative regulator of MO, NE, and NK cell infiltration upon LPS stimulation.

**Figure 2.**
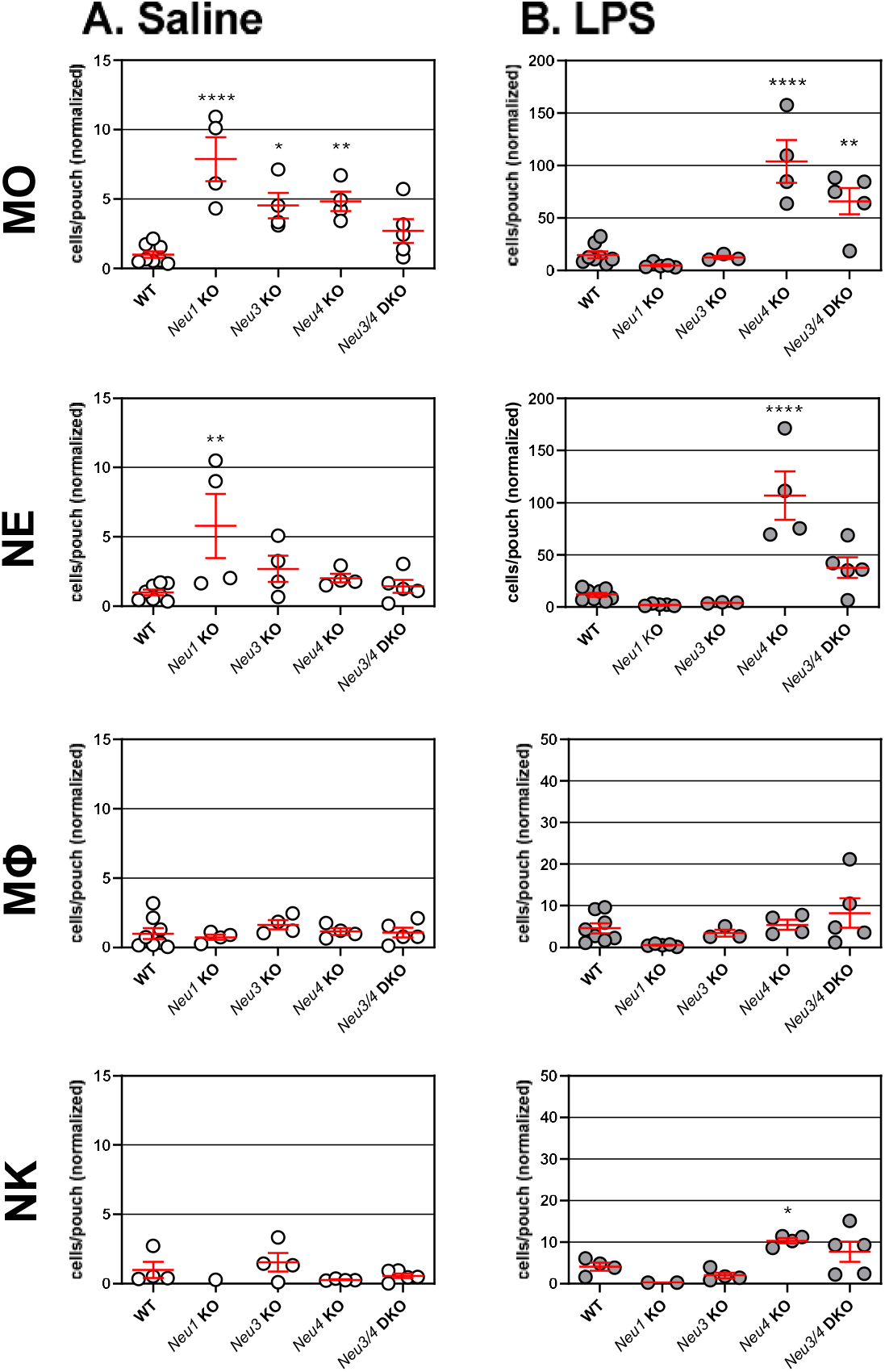
Changes in leukocyte populations in the air pouch model. Leukocytes collected from animals after saline (○), or LPS (●) treatment were counted by flow cytometry. The cells were identified after staining with marker-specific fluorochrome-conjugated antibodies as monocyte (MO), neutrophil (NE), macrophage (MΦ), or natural killer (NK) cells. Animals used were WT, *Neu1* KO, *Neu3* KO, *Neu4* KO and *Neu3/4* DKO mice. The air pouch exudate was collected after 9 h. The data are presented as cell counts (individual values and mean ± SEM) compared to the WT saline controls (monocyte, 7.4; neutrophil, 4.7; macrophage, 2.3; NK, 9.1 × 10^4^ cells/pouch). Means were compared to those of control groups using one-way ANOVA followed by a Dunnet’s t-test (*, p ≤ 0.05; **, p ≤ 0.01; ***, p ≤ 0.005; ****, p ≤ 0.0001). Raw cell counts are presented in **Figure SI1**.

### 3.3 Immunohistochemistry detects different levels of leukocyte infiltration in NEU-deficient animals

To add further support to our conclusions regarding leukocyte infiltration, tissue samples from the air pouch walls were analyzed for leukocyte infiltration (**Figure 3**, **Table SI2, Figure SI2**). Sections of the tissues were stained with H&E reagent and relative numbers of leukocytes in dermal and muscular layers were determined by microscopy. In the dermis layer of saline-treated *Neu1* KO animals, the levels of leukocytes were increased as compared to those in WT animals, but no difference was observed in the muscle layer. In contrast, in LPS-treated *Neu1* KO animals, leukocyte levels were significantly reduced in both dermis and muscle layers. In the muscle of LPS-treated *Neu3* KO animals, we observed a reduction in leukocytes. *Neu4* KO animals showed increased leukocyte counts in dermis and muscle layers with LPS treatment, and in muscle with saline treatment as compared with WT controls. This effect was attenuated in the *Neu3/4* DKO animals: while elevated counts in dermis and muscle were observed for saline controls, LPS treatment resulted in leukocyte counts either similar (dermis) or reduced (muscle) as compared with those in WT controls. These observations are consistent with results of cytometry analyses of serum and pouch exudates (vide supra) and provide additional support to the hypothesis that NEU1 and NEU3 act as positive regulators of infiltration, while NEU4 acts as a negative regulator.

**Figure 3.**
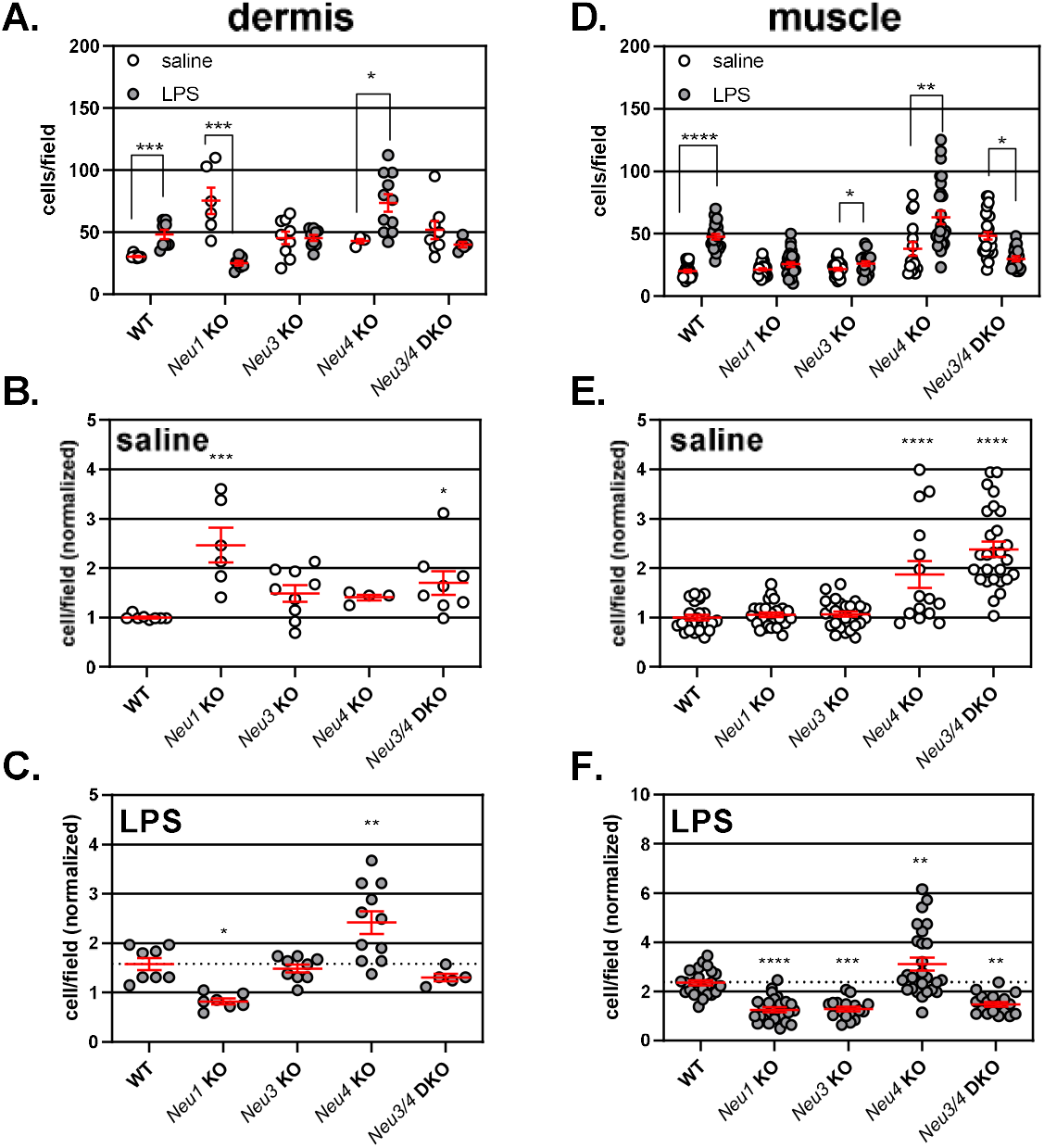
Levels of leukocytes in skin slices from the air pouch model. Tissue from the air pouch model was collected, sectioned, and stained (with H&E reagent). Regions of tissue were identified as dermis or muscle. See Fig SI2 for representative images of each region in different mice groups. Random fields from each region were used to determine leukocyte counts after saline (○) or LPS (●) treatment. Panels A. and D. show raw cell counts for each condition and tissue. Normalized cell counts after saline and LPS treatment are provided for dermis (B & C) and muscle (E & F) layers, respectively. The graphs show individual values and means ± SEM. Conditions were compared using a two-way ANOVA with Holm-Sidak’s multiple comparison (A and D) or one-way ANOVA followed by Dunnet’s multiple comparison (B, C, E, and F) (*, p ≤ 0.05; **, p ≤ 0.01; ***, p ≤ 0.005; ****, p ≤ 0.0001).

### 3.4 Effect of neuraminidases on cytokine levels in circulation and exudates

Based on the above findings, we concluded that NEU-deficient animals demonstrate altered regulation of leukocyte recruitment in the air-pouch model. One potential mechanism for these changes could involve the modulation of cytokine and chemokine production in the animals. To test this, air poach exudates and serum samples collected at sacrifice from WT, *Neu1* KO and *Neu4* KO mice were analyzed for a panel of 21 different cytokines and chemokines (**Table 2, Table SI3 & SI4**).

**Table 2:** Summary of changes to cytokine levels in *Neu1* and *Neu4* KO animals.

| Model |  | Basal activated† | Activated‡ | Attenuated§ |
| --- | --- | --- | --- | --- |
| *Neu1* KO | exudate | IL-10, IL-33 | G-CSF | G-CSF,<br>IL-1β, IL-6, IL-10,<br>MIP-1α, MIP-2,<br>TNF-α |
| | plasma | IL-1α, IL-1β,<br>IL-15, IL-25,<br>MIP-2 | TNF-α | G-CSF, RANTES,<br>IL-6, IL-10,<br>IFN-γ, MIP-1α, MIP-1β, MIP-2 |
| *Neu4* KO | exudate | | GM-CSF,<br>IL-1β,<br>IL-21, IL-33,<br>MIP-1α, MIP-1β, MIP-2 | G-CSF,<br>IFN-γ,<br>MCP-1 |
| | plasma | IP-10 | | G-CSF, RANTES,<br>IL-10,<br>MIP-1α, MIP-1β |

†, Basal activated was defined as having a *significant increase in saline for the model relative to saline control*.
‡, Activated was defined as having a *significant positive difference between control LPS and model LPS stimulation*.
§, Attenuated was defined as having a *significant negative difference between control LPS and model LPS stimulation*.

As expected, levels of pro-inflammatory cytokines and chemokines were low in the exudates of control (saline treated) WT mice and were drastically increased in the exudates of LPS-treated animals. The LPS-treated *Neu1* KO mice showed significantly attenuated levels of a range of inflammatory modulators such as G-CSF, IL-1β, IL-6, IL-10, MIP1-α, MIP-2, and TNF-α as compared with LPS-treated WT mice (**Figure 4, Figure SI3**). The *Neu1* KO animals also showed increased levels of GM-CSF in exudate after LPS treatment, while saline-treated animals had basal activation of IL-10 and IL-33. The LPS-treated *Neu4 KO* animals had distinct changes to exudate cytokine levels. We observed attenuation of G-CSF, IFN-γ, and MCP-1 and activation of GM-CSF, IL-1β, IL-21, IL-33, MIP1-α, MIP-1β, and MIP-2 in the exudates from *Neu4* KO animals.

**Figure 4.**
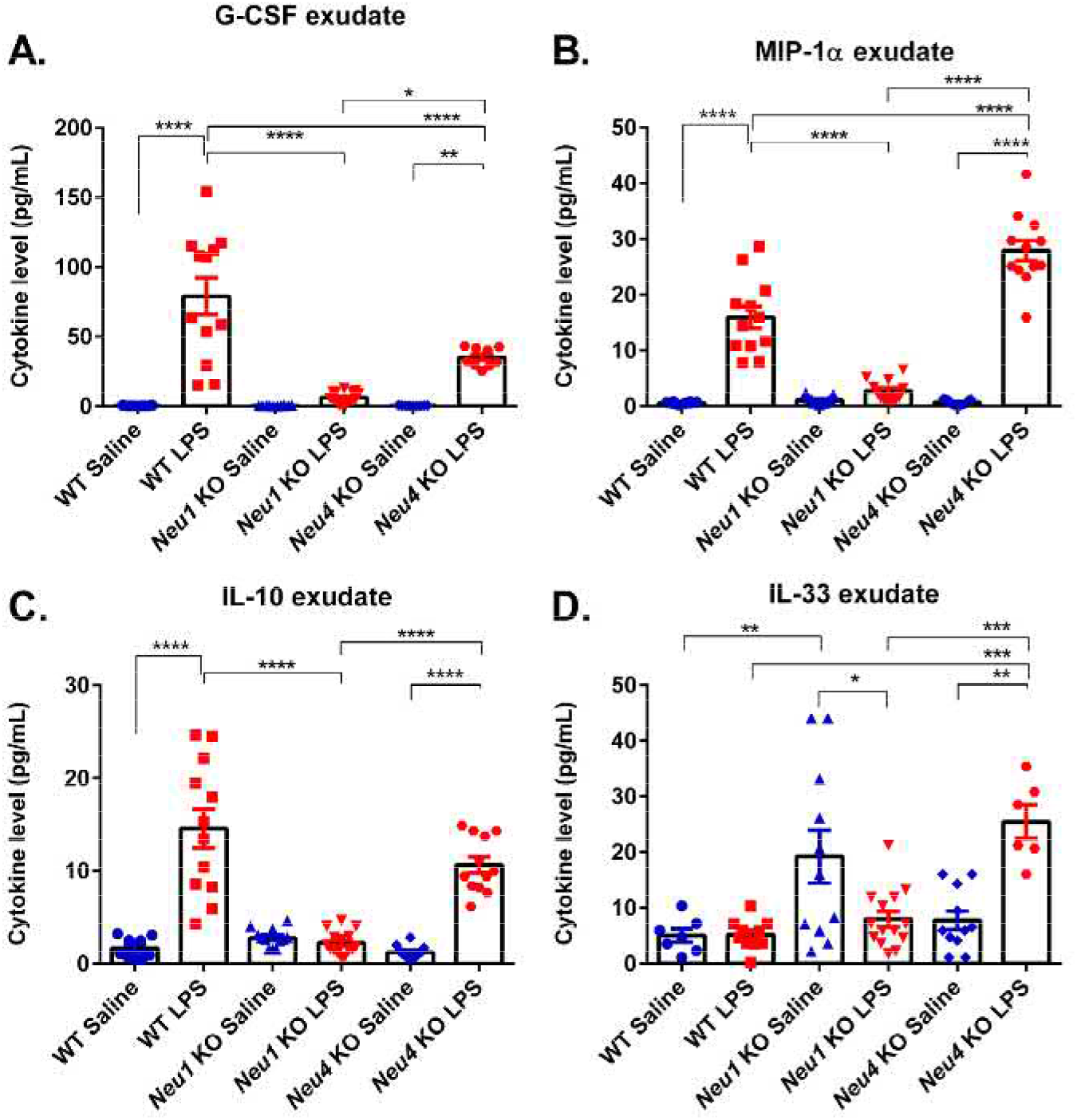
Cytokine levels in air poach exudates. Cytokines were analyzed using the Autoplex Analyser CS1000 system (Perkin Elmer, Waltham, MA) with a commercial ProcartaPlex Mouse Cytokine Panel Assay kit (Thermo Fisher Scientific Inc., Rockford, USA) in accordance with the manufacturer’s instruction. (**A**) G-CSF, (**B**) MIP-1α, (C) IL-10 and (D) IL-33 are shown as representatives, see Table 2 and Supporting information for other cytokines tested. Bars represent the mean values ± standard error of the mean (SEM) in units of pg/mL. Samples were compared to controls using a one-way ANOVA multiple comparison and Tukey’s test post-hoc analysis (*, p < 0.05; **, p < 0.01; ***, p < 0.001; ****, p < 0.0001).

We also examined plasma samples from these animals to observe systemic changes in cytokine levels. The LPS induction of 8 cytokines (G-CSF, RANTES, IL-6, IL-10, IFN-γ, MIP1-α, MIP-1β, and MIP-2) was significantly reduced in the plasma samples of *Neu1* KO as compared with LPS-treated WT mice, similar to exudate samples. The *Neu1* KO mice showed elevated levels of TNF-α in plasma. Cytokines significantly induced in saline-treated *Neu1* KO animals included IL-1α, IL-1β, IL-15, IL-25, and MIP-2 (**Figure 5**). The response of *Neu4* KO animals to LPS was attenuated for 5 cytokines in plasma (G-CSF, RANTES, IL-10, MIP-1α, and MIP-1β). Basal activation of IP-10 was observed in *Neu4* KO plasma samples.

**Figure 5.**
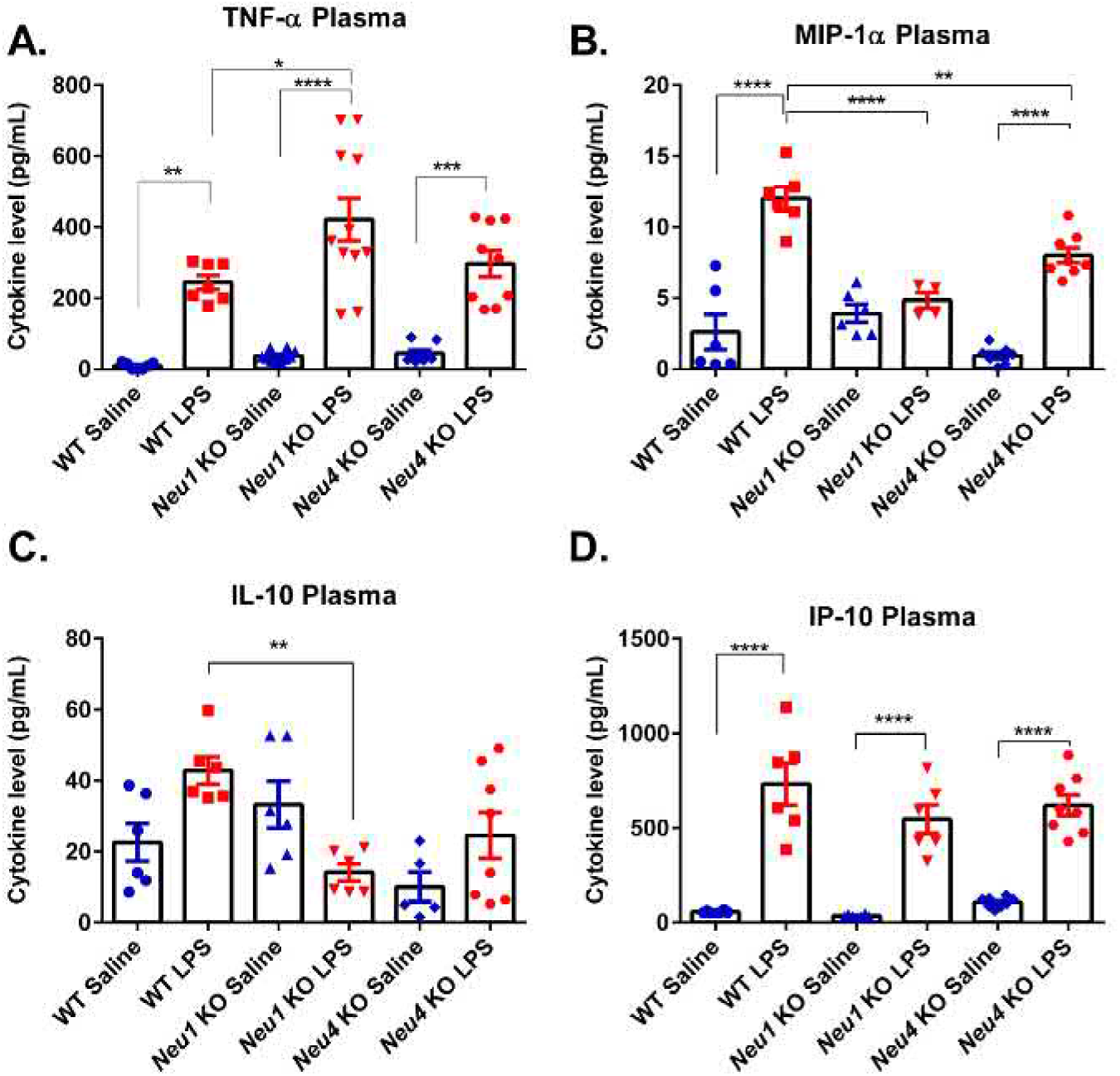
Cytokine levels in mouse plasma. Cytokines were analyzed using the Autoplex Analyser CS1000 system (Perkin Elmer, Waltham, MA) with a commercial ProcartaPlex Mouse Cytokine Panel Assay kit (Thermo Fisher Scientific Inc., Rockford, USA) in accordance with the manufacturer’s instruction. (**A**) TNF-α, (B) MIP-1α, (C) IL-10 and (D) IP-10 are shown as representatives, see Table 2 and Supporting information for other cytokines tested. Bars represent the mean values ± standard error of the mean (SEM) in units of pg/mL. Samples were compared to controls using a one-way ANOVA multiple comparison and Tukey’s test post-hoc analysis (*, p < 0.05; **, p < 0.01; ***, p < 0.001; ****, p < 0.0001)

### 3.5 Neuraminidase enzymes affect macrophage migration in vitro

Our observations from the air pouch model of inflammation clearly indicated that NEU isoenzymes can have dramatic impacts on leukocyte recruitment to sites of inflammation. To gain further insight into a potential mechanism, we investigated the effect of NEU deficiency on the migration of bone-marrow derived macrophages (BMDM) from these animals. For the migration assay, cultured BMDM from the indicated genotype were seeded into the top chamber of a transwell culture plate where the membrane separating the upper and lower chambers had been coated with fibronectin (FN). To the lower chamber of the plate, a chemoattractant (CCL2/MCP-1) was added. Nine hours after addition of BMDM to the upper chamber, we counted numbers of the leukocytes that infiltrated the membrane by microscopy (**Figure SI4**), the results are summarized in **Figure 6.** The migration experiment revealed that macrophages from *Neu1* KO animals were not able to migrate into a FN matrix. In contrast, the macrophages from *Neu3* KO animals demonstrated increased migration into the FN matrix. Macrophages from the *Neu4* KO animals showed no differences from WT controls.

**Figure 6.**
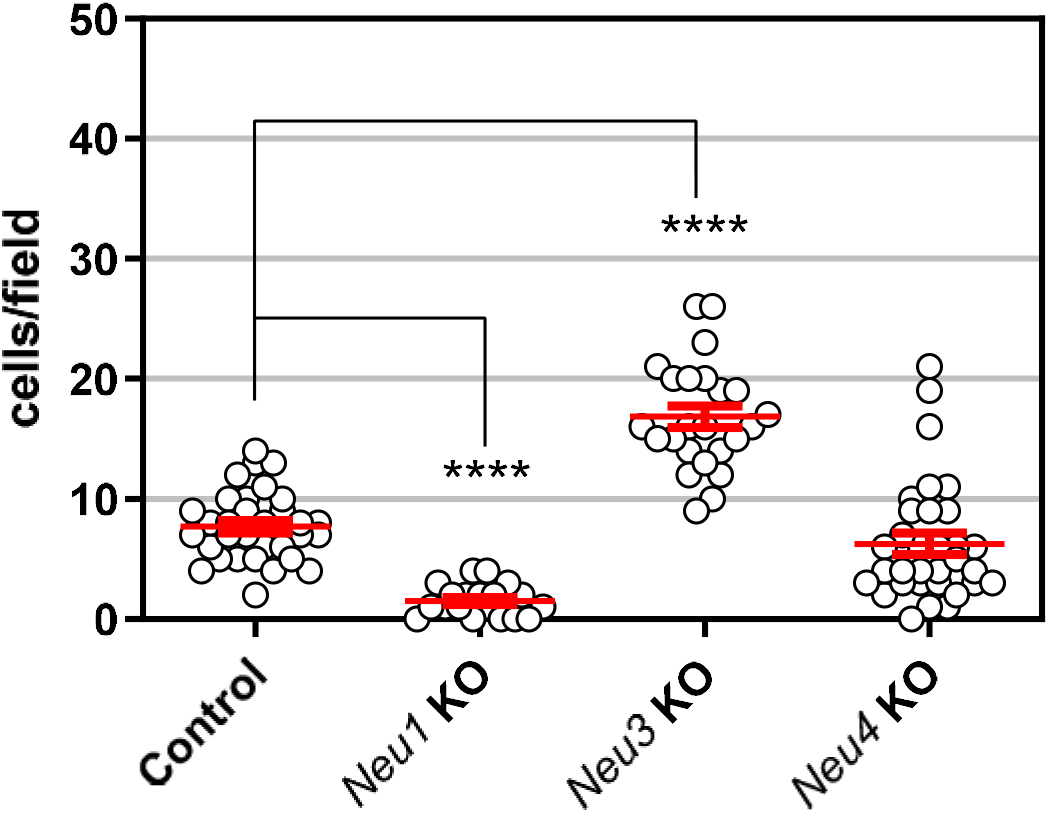
Migration of bone marrow-derived macrophages is affected by NEU expression. Macrophages were isolated and differentiated from bone marrow of WT, *Neu1* KO, *Neu3* KO, and *Neu4* KO mice. Cells (2.5 x10^4^) were placed in the upper chamber of the transmigration plate and the experiment was carried out for 5 h with MCP-1 as a chemoattractant in the lower chamber. The number of cells that infiltrated the FN-coated membrane was determined by counting of stained cells. Cell counts are plotted as mean ± SEM. Conditions were compared using a One-way ANOVA following Dunnett’s t-test for comparison between groups. (*, p ≤ 0.05; **, p ≤ 0.01; ***, p ≤ 0.005; ****, p ≤ 0.0001).

### 3.6 Effect of pharmacological inhibition of NEU on leukocyte subsets

Based on our findings, we considered that inhibitors of NEU isoenzymes should be able to recapitulate some of the effects of gene targeting on cell infiltration in the air-pouch inflammation model. We used previously reported inhibitors selective for NEU1, NEU3, and NEU4 (**Table 3**); compounds were dosed at 1 mg/kg body weight and delivered by IP injection 48 h, 24 h, and 9 h before sacrifice (**Table 1**, **Figure 7**).(Albohy *et al.*, 2013; Guo *et al.*, 2018a; Guo *et al.*, 2018b) We did not observe any significant effects for an inhibitor of NEU1 (**IN1**), which may indicate that the compound was not used at high enough dosage to counteract the most highly-expressed NEU isoenzyme. We observed an increase in leukocyte counts from exudate after treatment of WT animals with an inhibitor of NEU4 (**IN4**); however, no significant differences from controls were observed from inhibitor treatment in animals after LPS stimulation. We noted that a selective inhibitor of NEU3 (**IN3**) reduced the difference in leukocyte counts between saline and LPS treatments (**Figure 7A**). We examined the effects of NEU inhibitors on leukocyte subsets (**Figures SI5 & SI6; Table SI6)**. Both IN3 and IN4 treatments resulted in increased MO cell counts for saline treatment relative to control. Furthermore, IN3 treatment gave a significant reduction in MΦ counts after LPS treatment. We note that these experiments were at a single dosage and included a limited number of animals. As a result, follow up studies will be required to determine the potential of selective NEU inhibitors for affecting leukocyte infiltration.

**Figure 7.**
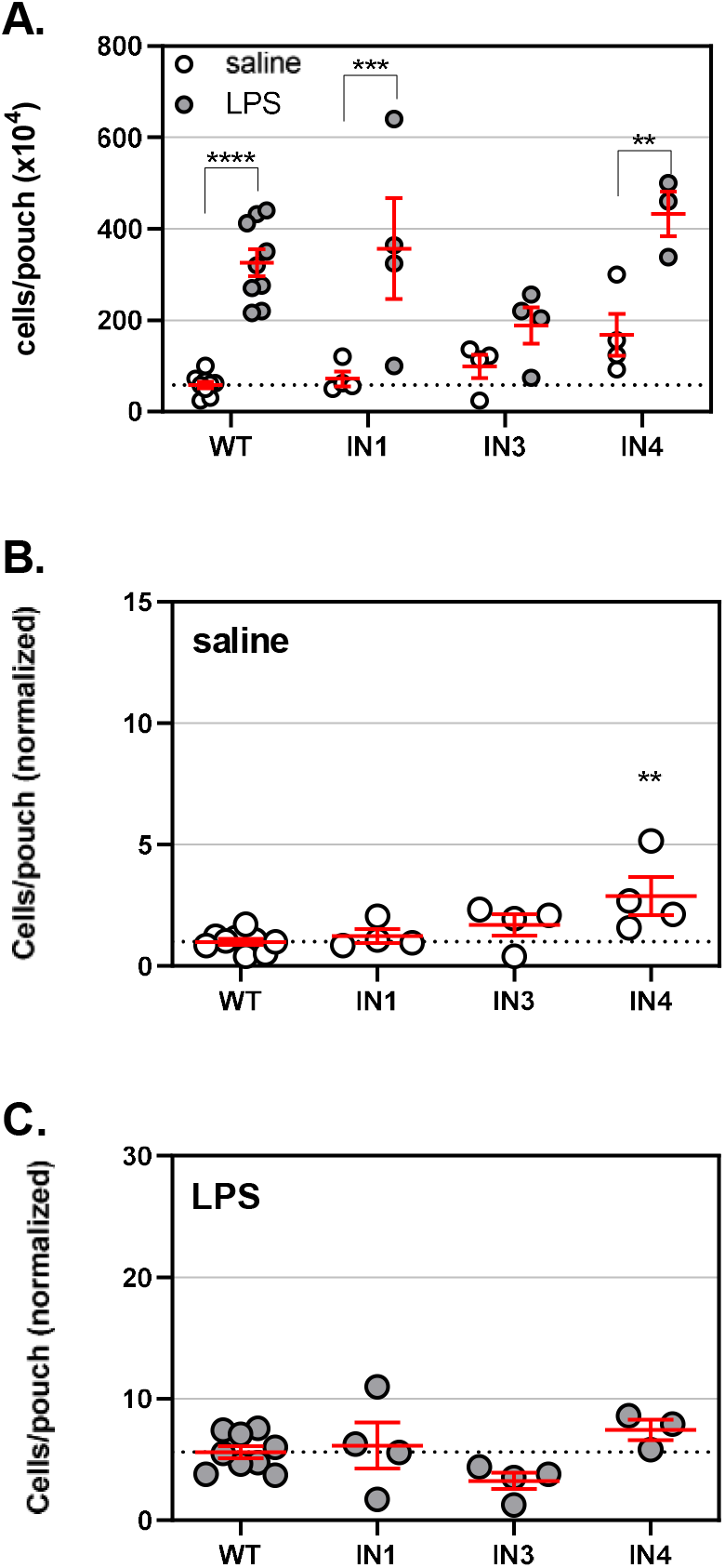
Leukocyte counts in the air pouch model are modulated by NEU inhibitors. An air pouch was formed in WT mice injected with inhibitor compounds or vehicle, followed by injection with saline or LPS to the air pouch and incubation for 9 h. Inhibitors that target NEU1 (**IN1**, CG14600), NEU3 (**IN3**, CG22600), and NEU4 (**IN4**, CY16600) were used. Mice were then sacrificed, the pouch exudate was collected, and cells were counted by flow cytometry. A. Total cell counts are presented for saline (○) and LPS (●) treatments. B. Cell counts for the saline treatment in the Neu KO mice are compared with those of WT mice. C. Cell counts for the LPS treatment are compared with those of WT mice. Data are presented as individual values and mean ± SEM. For panel A, comparisons were made using two-way ANOVA following Bonferroni multiple comparison t-test; for panel B & C, comparisons were made using one-way ANOVA with Dunnett’s t-test for multiple comparison (*, p ≤ 0.05; **, p ≤ 0.01; ***, p ≤ 0.005; ****, p ≤ 0.0001). Raw cell counts for leukocyte subsets are shown in **Figure SI5**.

**Table 3:**
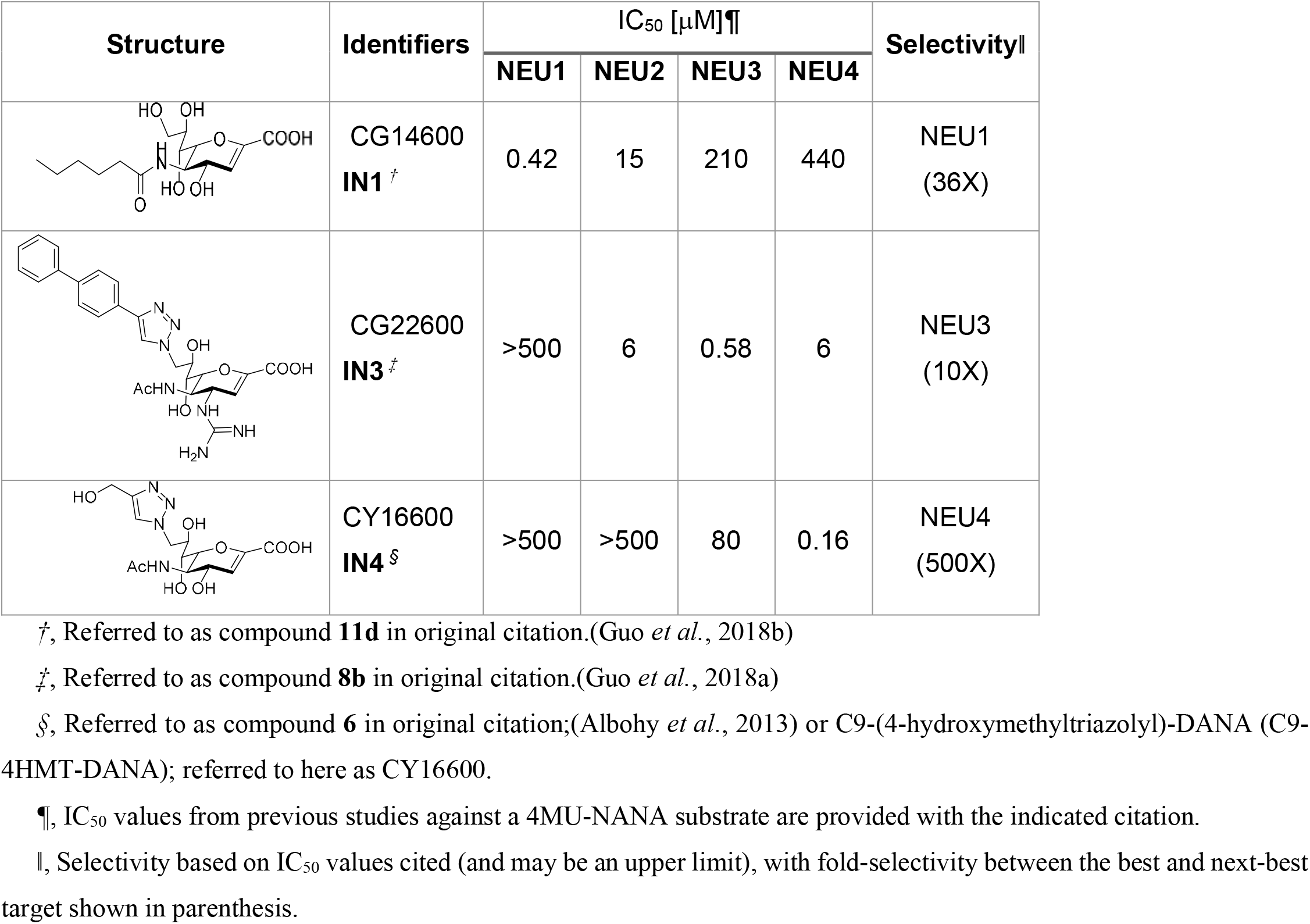
hNEU inhibitors used in this study.

| Structure | Identifiers | IC<sub>50</sub> [μM]<sup>¶</sup> |  |  |  | Selectivity |
| --- | --- | --- | --- | --- | --- | --- |
|  |  | NEU1 | NEU2 | NEU3 | NEU4 |  |
|  | CG14600<br>**IN1**<sup>‡</sup> | 0.42 | 15 | 210 | 440 | NEU1<br>(36X) |
|  | CG22600<br>**IN3**<sup>‡</sup> | >500 | 6 | 0.58 | 6 | NEU3<br>(10X) |
|  | CY16600<br>**IN4**<sup>§</sup> | >500 | >500 | 80 | 0.16 | NEU4<br>(500X) |

<sup>‡</sup>, Referred to as compound **11d** in original citation.(Guo *et al.*, 2018b)
<sup>‡</sup>, Referred to as compound **8b** in original citation.(Guo *et al.*, 2018a)
<sup>§</sup>, Referred to as compound **6** in original citation;(Albohy *et al.*, 2013) or C9-(4-hydroxymethyltriazolyl)-DANA (C9-4HMT-DANA); referred to here as CY16600.
<sup>¶</sup>, IC<sub>50</sub> values from previous studies against a 4MU-NANA substrate are provided with the indicated citation.
<sup>||</sup>, Selectivity based on IC<sub>50</sub> values cited (and may be an upper limit), with fold-selectivity between the best and next-best target shown in parenthesis.

## 4 Discussion

Using an air-pouch model of inflammation in gene-targeted mouse strains and a panel of specific neuraminidase inhibitors, we have investigated the involvement of endogenous neuraminidase enzymes in the migration and recruitment of leukocytes to the site of LPS-induced acute inflammation.(Muller, 2013) Our data show that NEU1 and NEU3 act as positive regulators of the recruitment of leukocytes to the site of inflammation. In contrast, NEU4 acts as a negative regulator of leukocyte recruitment. These results were confirmed by immunohistochemistry of dermal and muscular layers of the air pouch walls. Furthermore, LPS-treated *Neu1* KO, but not *Neu4* KO, animals have significantly reduced levels of proinflammatory cytokines IL-1β, MIP1-α, MIP-2, and TNF-α in air poach exudates. Thus, it can be concluded that NEU1 and NEU3, in general, exert pro-inflammatory effects, whereas NEU4 exerts anti-inflammatory effects.

Our data also demonstrated an increased basal level of inflammation in the control (saline treated) *Neu1* KO mice including higher levels of MO and NE in the air pouch washes, increased leukocyte infiltration in dermis layers of the pouch walls, and higher concentrations of circulating TNF-α, a pro-inflammatory cytokine. When interpreting these data, it is important to consider that complete deficiency of NEU1 results in a catabolic block in degradation of sialylated glycoproteins resulting in their lysosomal storage and leading to systemic metabolic disease, sialidosis.(Pshezhetsky & Ashmarina, 2001) Previous studies in human patients and mouse models of sialidosis have revealed that progressive neuroinflammation and leukocyte infiltration in peripheral tissues are the hallmarks of this disease.(Wu *et al*, 2010) Signs of neuroinflammation including microastrogliosis and increased brain levels of MIP-1α (CCL3) have been also reported for *Neu3* KO, *Neu4* KO, and *Neu3*/*Neu4* DKO mice but the levels were substantially lower as compared to *Neu1* KO animals.(Kho *et al*, 2020; Pan *et al.*, 2017)

Because the results obtained in KO mice suggested that pharmacological inhibition of NEU1 and NEU3 could be used for manipulating the inflammatory response, we further tested if specific inhibitors of neuraminidases were able to modulate leukocyte recruitment in the air pouch model. The results of our experiments with an inhibitor of NEU1 (**IN1**) were not conclusive due to the high variability between animals. However, treatment with an inhibitor of NEU3 (**IN3**) was consistent with the enzyme’s positive regulation of cell infiltration. Treatment with an inhibitor of NEU4 (**IN4**) was consistent with negative regulation of leukocyte recruitment. The inhibitors showed differential effects on leukocyte subsets, which may explain the divergence of these results from the gene-targeted animals. We observed that **IN3** reduced leukocyte recruitment in response to LPS in mice in similar fashion to the genetic inactivation of the *Neu3* gene. In contrast **IN4** increased leukocyte recruitment under basal conditions. Interestingly, both the NEU3 and NEU4 inhibitors increased basal levels of MO and NE in the pouch (saline treatment). While these data are generally supportive of our conclusion that NEU4 is a negative regulator while NEU1 and NEU3 are positive regulators of inflammation, the specific effects are different from those obtained by gene-targeting. It is important to also note that only a single dose and regimen for inhibitor administration were tested and the cohort size was limited. As a result, we cannot exclude that pharmacokinetics, bioavailability, or membrane permeability of the compounds may influence these conclusions. Further studies exploring different doses of inhibitors will be needed to clarify these effects.

A potential mechanism for regulation of leukocyte recruitment to the site of inflammation by NEU enzymes is modulation of pro-inflammatory cytokines and chemokine production. Indeed, our results indicate that NEU1 activity positively regulates cytokines directly implicated in the recruitment of leukocytes to the site of inflammation such as IL-1β, MIP-1α, and MIP-2.(Diab *et al*, 1999; Lotfi *et al*, 2019; Sherry *et al*, 1998) NEU1 overexpressing MΦ are known to have increased expression of IL-1β and TNF-α, consistent with our observation of these cytokines being attenuated in the *Neu1* KO model.(Sieve *et al*, 2018) Previous studies have shown that treatment of cells with exogenous bacterial neuraminidases exert differential effects on cytokines and chemokines. For example, Stamatos and coworkers showed that exogenous treatment of purified human MO with neuraminidase from *Clostridium perfringens* increased production of IL-6, MIP-1α (CCL3) and MIP-1β (CCL4), but had no effect on RANTES (CCL5), IL-10, TNF-α, IFN-γ, and IL-1β.(Stamatos *et al*, 2004) The differences in the results on the production of different cytokines and chemokines between this and our study could be due to different experimental approaches and enzyme specificities.

*Neu4* KO mice, in contrast to *Neu1* KO mice, had fewer changes to cytokine profiles. Our data regarding cell migration to the air pouch indicated elevated accumulation of MO, NE, and NK cells in the *Neu4* KO model. This observation may be partly explained by elevated cytokine levels observed in exudate samples from these animals. We observed activation of 7 cytokines from *Neu4* KO exudate samples. Among these are cytokines that are known to act as chemoattractants for leukocytes including MO, NE, or NK cells.(Choi *et al*, 2009; Diab *et al.*, 1999; Gomez-Cambronero *et al*, 2003; Lotfi *et al.*, 2019; Pelletier *et al*, 2004; Sherry *et al.*, 1998) Together, these data suggest that NEU4 and NEU1 modulate cytokine levels which may explain the changes in leukocyte migration observed in these models. Further experiments will be required to test if cytokine levels are directly or indirectly affected by NEU deficiencies; however, our current data provide evidence for clear differences between WT, *Neu1* KO and *Neu4* KO mice cytokine levels and warrant future investigation.

We have demonstrated earlier that levels of NEU1 are increased at least 10-fold during the differentiation of MO to MΦ, and that the enzyme is targeted from lysosomes to the cell membrane.(Liang *et al*, 2006; Stamatos *et al*, 2005) Furthermore, increased sialylation of cell surface receptors targeted by NEU1, including Fc-γ receptors on MΦ, have been demonstrated in *CathA^S190Aneo^* mice with ~90% reduction of NEU1 activity.(Pshezhetsky & Hinek, 2011; Seyrantepe *et al.*, 2010) The *Neu1* KO mouse is completely deficient in NEU1 and is likely to show enhanced sialylation of NEU1-targeted glycoconjugates including receptors and proteins involved in signalling cascades activated by LPS. In our study, we used LPS, a major component of the cell wall of Gram-negative bacteria, to induce inflammation in the air pouch. The binding of LPS to TLR-4 via CD14 (a GPI-anchored homodimer) induces TLR-4 dimerization and activates a signalling cascade that culminates in the activation of NF-kB, and production of pro-inflammatory cytokines and chemokines.(Kawasaki & Kawai, 2014) We have reported that the TLR-4-induced signaling causes activation and translocation of endogenous NEU1 to the cell surface. The activated NEU1 desialylates TLR-4, and this process plays a critical role in the TLR-4-mediated activation of NF-kB.(Amith *et al.*, 2010; Pshezhetsky & Ashmarina, 2013) This mechanism may explain the reduced production of pro-inflammatory cytokines and chemokines in the *Neu1* KO mouse. Not surprisingly, neuraminidase potentiates LPS-induced acute lung injury in mice.(Feng *et al*, 2013) *Neu1* KO has been found to block LPS induction of TNF-α and IL-1β in monocytes.(Sieve *et al.*, 2018) Besides potentiating TLR-4 signalling, cell surface protein hypersialylation may affect production of cytokines and chemokines by other mechanisms such as masking of galectin receptors or engagement of inhibitory Siglecs by trans- and cis-acting sialylated epitopes.(Pearce & Läubli, 2016; Schauer, 2009) Further studies are required to clarify specific links between NEU1 and inflammasome activation.

The rolling and tethering of leukocytes onto endothelial cells is a first step in the extravasation of leukocytes to the sites of inflammation in the body.(Muller, 2013) The sialylated epitopes sLe^a^ and sLe^x^ (CD15s) are present on a variety of glycoproteins, such as PSGL-1 and CD44, and are critical for extravasation of normal leukocytes. These sialylated epitopes are also abundantly expressed on cancer cells and mediate their interaction through selectins on endothelial cells. Interestingly, Neu4S, the short cytoplasmic NEU4 isoform expressed in peripheral tissues, is downregulated in cancer and can desialylate these binding epitopes.(Shiozaki *et al*, 2011) This may suggest a molecular mechanism by which NEU4 could act as a negative regulator of leukocyte recruitment to a site of inflammation.

Our study revealed a positive role for NEU3 in the recruitment of leukocytes. Previous studies have found that neuraminidases can affect leukocyte recruitment by modulating activities of adhesion molecules.(Wright & Cooper, 2014) Our previous work has suggested that NEU3 mediates desialylation of gangliosides and glycoproteins which may inhibit cis-interactions with β1 integrins (α4β1 and α5β1; also known as VLA4 and VLA5, respectively).(Howlader *et al.*, 2019; Jia *et al*, 2016) Here, we confirmed that BMDM from *Neu3* KO animals had increased in vitro chemotaxis through a FN-coated membrane, consistent with this isoenzyme acting as a negative regulator of cell migration in response to a chemoattractant (MCP-1/CCL2). In contrast, *Neu1* KO animals showed reduced migration in vitro. Further studies are required to clarify the mechanism by which NEU1 induces leukocyte transmigration, but it is notable that the MCP-1 receptor, CCR2, is sialylated.(Matsubara *et al*, 2015) BMDM from *Neu4* KO mice did not show differences from control in the migration assay, which may indicate that the enzyme regulates a different aspect of leukocyte recruitment. We hope that our findings will spur future investigations of leukocyte activation to explore the specific mechanisms responsible for these effects more fully.

In conclusion, this study has demonstrated that neuraminidase isoenzymes play a key role in LPS-stimulated acute inflammatory response. These effects were highly dependent upon the NEU isoenzyme tested. We found that NEU1 and NEU3 activity was pro-inflammatory, while NEU4 activity was anti-inflammatory. Together, these results bolster the case for differential involvement of NEU isoenzymes in the activation and infiltration of leukocytes during inflammation. This discovery could provide new targets for therapeutics and may indicate an unrecognized role for NEU in immune cell regulation. Further work is necessary to determine the specific mechanisms regulated by each enzyme tested here.

## Supporting information

Additional Supporting information can be found online in the Supporting Information section.

## Acknowledgements

We are grateful for the use of facilities provided by the University of Alberta Department of Chemistry. IHC was performed by Sullen Lamb, Histology Lab Services, University of Alberta.

## Author contributions

MAH, ED, SS, AA, AVP, and CWC designed research; MAH, ED, and SS performed research and analyzed data; TG contributed new reagents; MAH, AA, AVP, and CWC wrote the manuscript.

## Abbreviations

BW: body weight
BMDM: bone marrow-derived macrophage
DKO: double knock out
IN1: inhibitor of NEU1 (CG14600)
IN3: inhibitor of NEU3 (CG22600)
IN4: inhibitor of NEU4 (CY16600)
KO: knock out
LPS: lipopolysaccharide
MO: monocytes
MΦ: macrophages
NE: neutrophil
NEU1: neuraminidase 1
NEU3: neuraminidase 3
NEU4: neuraminidase 4
NK: natural killer cells
WT: wild type

