## Supporting information for "The Janus-like role of neuraminidase isoenzymes in inflammation"

**Table SI1. Leukocyte subset counts in the air pouch model.**

| MONOCYTES | Saline (cells x$10^4$) | | LPS (cells x$10^4$) | |
|---|---|---|---|---|
| | Mean ± SEM | N | Mean ± SEM | N |
| **WT C57BL6** | 7 ± 2 | 8 | 110 ± 24 | 8 |
| **NEU3 4 DKO** | 20 ± 6 | 5 | 490 ± 94 | 5 |
| **NEU 4 KO** | 36 ± 5 | 4 | 771 ± 150 | 4 |
| **NEU 3 KO** | 34 ± 7 | 4 | 91 ± 12 | 3 |
| **NEU1 KO** | 59 ± 12 | 4 | 35 ± 7 | 5 |

| NEUTROPHILS | Saline (cells x$10^4$) | | LPS (cells x$10^4$) | |
|---|---|---|---|---|
| | Mean ± SEM | N | Mean ± SEM | N |
| **WT C57BL6** | 5 ± 1 | 8 | 57 ± 9 | 8 |
| **NEU3 4 DKO** | 7 ± 2 | 5 | 179 ± 47 | 5 |
| **NEU 4 KO** | 10 ± 1 | 4 | 507 ± 111 | 4 |
| **NEU 3 KO** | 13 ± 5 | 4 | 20 ± 2 | 3 |
| **NEU1 KO** | 28 ± 11 | 4 | 10 ± 2 | 5 |

| MACROPHAGES | Saline (cells x$10^4$) | | LPS (cells x$10^4$) | |
|---|---|---|---|---|
| | Mean ± SEM | N | Mean ± SEM | N |
| **WT C57BL6** | 2 ± 1 | 8 | 11 ± 3 | 8 |
| **NEU3 4 DKO** | 3 ± 1 | 5 | 20 ± 8 | 5 |
| **NEU 4 KO** | 3 ± 1 | 4 | 13 ± 3 | 4 |
| **NEU 3 KO** | 4 ± 1 | 4 | 8 ± 2 | 3 |
| **NEU1 KO** | 2 ± 1 | 4 | 1 ± 1 | 5 |

| NK CELLS | Saline (cells x$10^4$) | | LPS (cells x$10^4$) | |
|---|---|---|---|---|
| | Mean ± SEM | N | Mean ± SEM | N |
| **WT C57BL6** | 9 ± 5 | 4 | 38 ± 9 | 4 |
| **NEU3 4 DKO** | 5 ± 2 | 5 | 70 ± 22 | 5 |
| **NEU 4 KO** | 3 ± 1 | 4 | 95 ± 6 | 4 |
| **NEU 3 KO** | 14 ± 6 | 4 | 18 ± 6 | 4 |
| **NEU1 KO** | 1 ± 1 | 2 | 2 ± 0.1 | 2 |

| B Cells | Saline (cells x$10^4$) | | LPS (cells x$10^4$) | |
|---|---|---|---|---|
| | Mean ± SEM | N | Mean ± SEM | N |
| **WT C57BL6** | 2 ± 1 | 4 | 15 ± 7 | 4 |
| **NEU3 4 DKO** | 4 ± 1 | 5 | 94 ± 60 | 5 |
| **NEU 4 KO** | 4 ± 1 | 4 | 3 ± 1 | 4 |
| **NEU 3 KO** | 2 ± 1 | 4 | 6 ± 2 | 4 |
| **NEU1 KO** | 3 ± 3 | 2 | 1 ± 0.1 | 2 |

| T CELLS | Saline (cells x$10^4$) | | LPS (cells x$10^4$) | |
|---|---|---|---|---|
| | Mean ± SEM | N | Mean ± SEM | N |
| **WT C57BL6** | 5 ± 4 | 4 | 30 ± 20 | 4 |
| **NEU3 4 DKO** | 5 ± 1 | 5 | 154 ± 46 | 5 |
| **NEU 4 KO** | 5 ± 1 | 4 | 34 ± 10 | 4 |
| **NEU 3 KO** | 12 ± 3 | 4 | 34 ± 8 | 4 |
| **NEU1 KO** | 2 ± 2 | 2 | 4 ± 1 | 2 |

Results expressed as mean ± standard error of the mean (SEM).

### Table SI2. Leukocyte counts in skin slices from the air pouch model.

| Muscle layer | Saline (cells/field) | | LPS (cells/field) | |
|---|---|---|---|---|
| | Mean ± SEM | N | Mean ± SEM | N |
| **WT C57BL6** | 20 ± 1 | 25 | 48 ± 3 | 24 |
| **NEU1 KO** | 21 ± 1 | 24 | 26 ± 2 | 26 |
| **NEU3 KO** | 22 ± 1 | 27 | 26 ± 2 | 17 |
| **NEU4 KO** | 38 ± 6 | 15 | 63 ± 5 | 27 |
| **NEU3/4 DKO** | 48 ± 3 | 27 | 30 ± 2 | 18 |

| Dermis layer | Saline (/field) | | LPS (/field) | |
|---|---|---|---|---|
| | Mean ± SEM | N | Mean ± SEM | N |
| **WT C57BL6** | 31 ± 1 | 8 | 48 ± 4 | 8 |
| **NEU1 KO** | 75 ± 11 | 6 | 25 ± 2 | 7 |
| **NEU3 KO** | 45 ± 5 | 9 | 45 ± 2 | 9 |
| **NEU4 KO** | 43 ± 2 | 4 | 74 ± 7 | 11 |
| **NEU3/4 DKO** | 52 ± 7 | 8 | 40 ± 2 | 5 |

**Table SI3. Serum cytokine levels from plasma samples**

| Plasma Cytokine Profile [1] | | | | | | | | | | | | | | | | | | |
|---|---|---|---|---|---|---|---|---|---|---|---|---|---|---|---|---|---|---|
| | **WT Saline** | | | **WT LPS** | | | **NEU1 KO Saline** | | | **NEU1 KO LPS** | | | **NEU4 KO Saline** | | | **NEU4 KO LPS** | | |
| | Mean | ± | SEM | Mean | ± | SEM | Mean | ± | SEM | Mean | ± | SEM | Mean | ± | SEM | Mean | ± | SEM |
| **G CSF** | 5 | ± | 1 | 672 | ± | 97 | 28 | ± | 2 | 95 | ± | 26 | 11 | ± | 3 | 330 | ± | 39 |
| **IL-1 alpha** | 13 | ± | 3 | 41 | ± | 4 | 49 | ± | 7 | 18 | ± | 9 | 9 | ± | 3 | 33 | ± | 11 |
| **IL-1 beta** | 1 | ± | 0.1 | 7 | ± | 1 | 7 | ± | 1 | 3 | ± | 0.9 | 2 | ± | 0.8 | 7 | ± | 2 |
| **IL-2** | 3 | ± | 1 | 63 | ± | 14 | 57 | ± | 26 | 51 | ± | 13 | 73 | ± | 24 | 123 | ± | 26 |
| **IL-6** | 19 | ± | 7 | 557 | ± | 92 | 63 | ± | 5 | 181 | ± | 80 | 61 | ± | 9 | 407 | ± | 23 |
| **GM CSF** | 8 | ± | 2 | 35 | ± | 5 | 34 | ± | 4 | 18 | ± | 5 | 14 | ± | 4 | 31 | ± | 9 |
| **IL-25 IL-17** | 10 | ± | 2 | 181 | ± | 27 | 163 | ± | 15 | 65 | ± | 24 | 61 | ± | 23 | 152 | ± | 50 |
| **IL-10** | 23 | ± | 5 | 43 | ± | 4 | 33 | ± | 7 | 14 | ± | 2 | 10 | ± | 4 | 25 | ± | 6 |
| **IL-21** | 3 | ± | 0.7 | 21 | ± | 5 | 29 | ± | 5 | 12 | ± | 4 | 5 | ± | 1 | 33 | ± | 12 |
| **IP-10** | 57 | ± | 3 | 732 | ± | 111 | 35 | ± | 3 | 546 | ± | 75 | 107 | ± | 10 | 620 | ± | 55 |
| **IL-15/ IL-15R** | 3 | ± | 0.5 | 23 | ± | 3 | 21 | ± | 2 | 11 | ± | 2 | 8 | ± | 2 | 23 | ± | 6 |
| **RANTES** | 16 | ± | 2 | 924 | ± | 144 | 54 | ± | 9 | 414 | ± | 114 | 13 | ± | 4 | 362 | ± | 21 |
| **MCP-1 CCL2** | 75 | ± | 0 | 2473 | ± | 306 | 67 | ± | 24 | 1213 | ± | 357 | 35 | ± | 6 | 947 | ± | 82 |
| **MIP-1 alpha** | 3 | ± | 1 | 12 | ± | 0.8 | 4 | ± | 0.6 | 5 | ± | 0.6 | 0.9 | ± | 0.2 | 8 | ± | 0.5 |
| **MIP-1 beta** | 2 | ± | 0.3 | 107 | ± | 10 | 7 | ± | 1 | 35 | ± | 2 | 3 | ± | 0.6 | 51 | ± | 7 |
| **MIP-2** | 4 | ± | 0.2 | 17 | ± | 2 | 14 | ± | 2 | 10 | ± | 2 | 6 | ± | 1 | 14 | ± | 2 |
| **INF-γ** | 2 | ± | 0.8 | 5 | ± | 0.7 | 4 | ± | 0.7 | 2 | ± | 0.6 | 0.4 | ± | 0.2 | 4 | ± | 1 |
| **TNF-α** | 9 | ± | 3 | 244 | ± | 20 | 37 | ± | 5 | 421 | ± | 60 | 44 | ± | 10 | 297 | ± | 37 |
| **IL-18** | 1092 | ± | 166 | 1970 | ± | 162 | 943 | ± | 170 | 1456 | ± | 160 | 1275 | ± | 28 | 1356 | ± | 131 |
| **IL-33** | 18 | ± | 2 | 327 | ± | 81 | 96 | ± | 29 | 312 | ± | 68 | 17 | ± | 3 | 452 | ± | 178 |
| **IL-12p40** | 37 | ± | 7 | 49 | ± | 6 | 42 | ± | 6 | 167 | ± | 79 | 24 | ± | 3 | 46 | ± | 2 |

1. Results expressed as mean ± standard error of the mean (SEM); all values are in pg/mL.

### Table SI4. Serum cytokine levels from exudate

| Exudate Cytokine Profile [1] | | | | | | | | | | | | | | | | | | |
|---|---|---|---|---|---|---|---|---|---|---|---|---|---|---|---|---|---|---|
| | WT Saline | | | WT LPS | | | NEU1 KO Saline | | | NEU1 KO LPS | | | NEU4 KO Saline | | | NEU4 KO LPS | | |
| | Mean | ± | SEM | Mean | ± | SEM | Mean | ± | SEM | Mean | ± | SEM | Mean | ± | SEM | Mean | ± | SEM |
| **G CSF** | 0.3 | ± | 0.06 | 79 | ± | 13 | 0.3 | ± | 0.06 | 6 | ± | 1 | 0.3 | ± | 0.08 | 35 | ± | 2 |
| **IL-1 alpha** | 10 | ± | 1 | 16 | ± | 1 | 13 | ± | 1 | 12 | ± | 1 | 5 | ± | 1 | 14 | ± | 1 |
| **IL-1 beta** | 0.6 | ± | 0.2 | 5 | ± | 0.5 | 0.2 | ± | 0.06 | 1 | ± | 0.2 | 0.2 | ± | 0.03 | 7 | ± | 0.4 |
| **IL-2** | 5 | ± | 0.8 | 6 | ± | 0.4 | 7 | ± | 0.7 | 6 | ± | 0.8 | 6 | ± | 2 | 7 | ± | 1 |
| **IL-6** | 269 | ± | 118 | 1050 | ± | 213 | 4 | ± | 1 | 140 | ± | 48 | 13 | ± | 3 | 840 | ± | 139 |
| **GM CSF** | 1 | ± | 0.3 | 2 | ± | 0.2 | 2 | ± | 0.5 | 2 | ± | 0.3 | 0.6 | ± | 0.1 | 4 | ± | 0.07 |
| **IL-25 IL-17** | 12 | ± | 2 | 16 | ± | 3 | 16 | ± | 2 | 17 | ± | 5 | 6 | ± | 1 | 23 | ± | 3 |
| **IL-10** | 2 | ± | 0.4 | 15 | ± | 2 | 3 | ± | 0.3 | 2 | ± | 0.3 | 1 | ± | 0.2 | 11 | ± | 0.9 |
| **IL-21** | 6 | ± | 0.3 | 3 | ± | 0.6 | 6 | ± | 0.8 | 6 | ± | 0.7 | 5 | ± | 0.5 | 5 | ± | 0.5 |
| **IP-10** | 10 | ± | 0.4 | 409 | ± | 62 | 14 | ± | 2 | 268 | ± | 29 | 23 | ± | 5 | 409 | ± | 53 |
| **IL-15/ IL-15R** | 0.7 | ± | 0.1 | 3 | ± | 0.3 | 0.9 | ± | 0.2 | 1 | ± | 0.1 | 0.9 | ± | 0.1 | 3 | ± | 0.3 |
| **RANTES** | 5 | ± | 0.8 | 190 | ± | 31 | 6 | ± | 0.6 | 179 | ± | 25 | 3 | ± | 0.3 | 184 | ± | 22 |
| **MCP-1 CCL2** | 196 | ± | 36 | 4445 | ± | 1054 | 55 | ± | 8 | 3383 | ± | 535 | 70 | ± | 5 | 1705 | ± | 260 |
| **MIP-1 alpha** | 0.6 | ± | 0.06 | 16 | ± | 2 | 1 | ± | 0.2 | 3 | ± | 0.6 | 0.7 | ± | 0.1 | 28 | ± | 2 |
| **MIP-1 beta** | 0.7 | ± | 0.1 | 57 | ± | 9 | 0.9 | ± | 0.1 | 17 | ± | 4 | 2 | ± | 0.1 | 83 | ± | 5 |
| **MIP-2** | 16 | ± | 6 | 21 | ± | 2 | 7 | ± | 1 | 8 | ± | 0.4 | 8 | ± | 0.8 | 33 | ± | 3 |
| **INF-γ** | 0.08 | ± | 0.03 | 7 | ± | 0.7 | 0.1 | ± | 0.03 | 1 | ± | 0.2 | 0.05 | ± | 0.02 | 4 | ± | 0.7 |
| **TNF-α** | 2 | ± | 0.5 | 16 | ± | 2 | 3 | ± | 0.3 | 5 | ± | 0.6 | 3 | ± | 0.4 | 15 | ± | 2 |
| **IL-18** | 6 | ± | 0.8 | 19 | ± | 3 | 15 | ± | 2 | 17 | ± | 3 | 4 | ± | 0.8 | 25 | ± | 2 |
| **IL-33** | 5 | ± | 1 | 5 | ± | 1 | 19 | ± | 5 | 8 | ± | 1 | 8 | ± | 2 | 25 | ± | 345 |
| **IL-12p40** | 45 | ± | 8 | 208 | ± | 38 | 27 | ± | 4 | 154 | ± | 31 | 23 | ± | 3 | 167 | ± | 17 |

1. Results expressed as mean ± standard error of the mean (SEM); all values are in pg/mL.

### Table SI5. Transmigration of macrophages isolated from different genotypic mice.

| | (Cells/field) | |
|---|---|---|
| | Mean ± SEM | N |
| **WT C57BL6** | 8 ± 1 | 32 |
| **NEU1 KO** | 2 ± 1 | 19 |
| **NEU3 KO** | 17 ± 1 | 24 |
| **NEU4 KO** | 6 ± 1 | 32 |

### Table SI6. Leukocyte populations in the air pouch model after inhibitor treatment.

| **MONOCYTES** | **Saline (cells x10$^4$)** | | **LPS (cells x10$^4$)** | |
|---|---|---|---|---|
| | Mean ± SEM | N | Mean ± SEM | N |
| **WT C57BL6** | 7 ± 2 | 8 | 75 ± 8 | 8 |
| **IN1** | 17 ± 4 | 4 | 157 ± 48 | 5 |
| **IN3** | 37 ± 13 | 4 | 78 ± 22 | 4 |
| **IN4** | 25 ± 1 | 3 | 182 ± 46 | 3 |

| **NEUTROPHILS** | **Saline (cells x10$^4$)** | | **LPS (cells x10$^4$)** | |
|---|---|---|---|---|
| | Mean ± SEM | N | Mean ± SEM | N |
| **WT C57BL6** | 5 ± 1 | 8 | 57 ± 9 | 8 |
| **IN1** | 7 ± 1 | 4 | 121 ± 56 | 4 |
| **IN3** | 26 ± 7 | 4 | 42 ± 9 | 4 |
| **IN4** | 22 ± 7 | 4 | 146 ± 41 | 3 |

| **MACROPHAGES** | **Saline (cells x10$^4$)** | | **LPS (cells x10$^4$)** | |
|---|---|---|---|---|
| | Mean ± SEM | N | Mean ± SEM | N |
| **WT C57BL6** | 2 ± 1 | 8 | 11 ± 3 | 8 |
| **IN1** | 2 ± 1 | 5 | 2 ± 1 | 5 |
| **IN3** | 2 ± 1 | 4 | 1 ± .2 | 4 |
| **IN4** | 2 ± 1 | 4 | 5 ± 2 | 3 |

| **NK CELLS** | **Saline (cells x10$^4$)** | | **LPS (cells x10$^4$)** | |
|---|---|---|---|---|
| | Mean ± SEM | (N) | Mean ± SEM | (N) |
| **WT C57BL6** | 4 ± 1 | (3) | 38 ± 9 | (4) |
| **IN1** | 13 ± 2 | (4) | 24 ± 8 | (4) |
| **IN3** | 3 ± 1 | (4) | 7 ± 1 | (4) |
| **IN4** | 30 ± 10 | (4) | 65 ± 31 | (3) |

Number of cells found in the air pouch was counted by FACS.

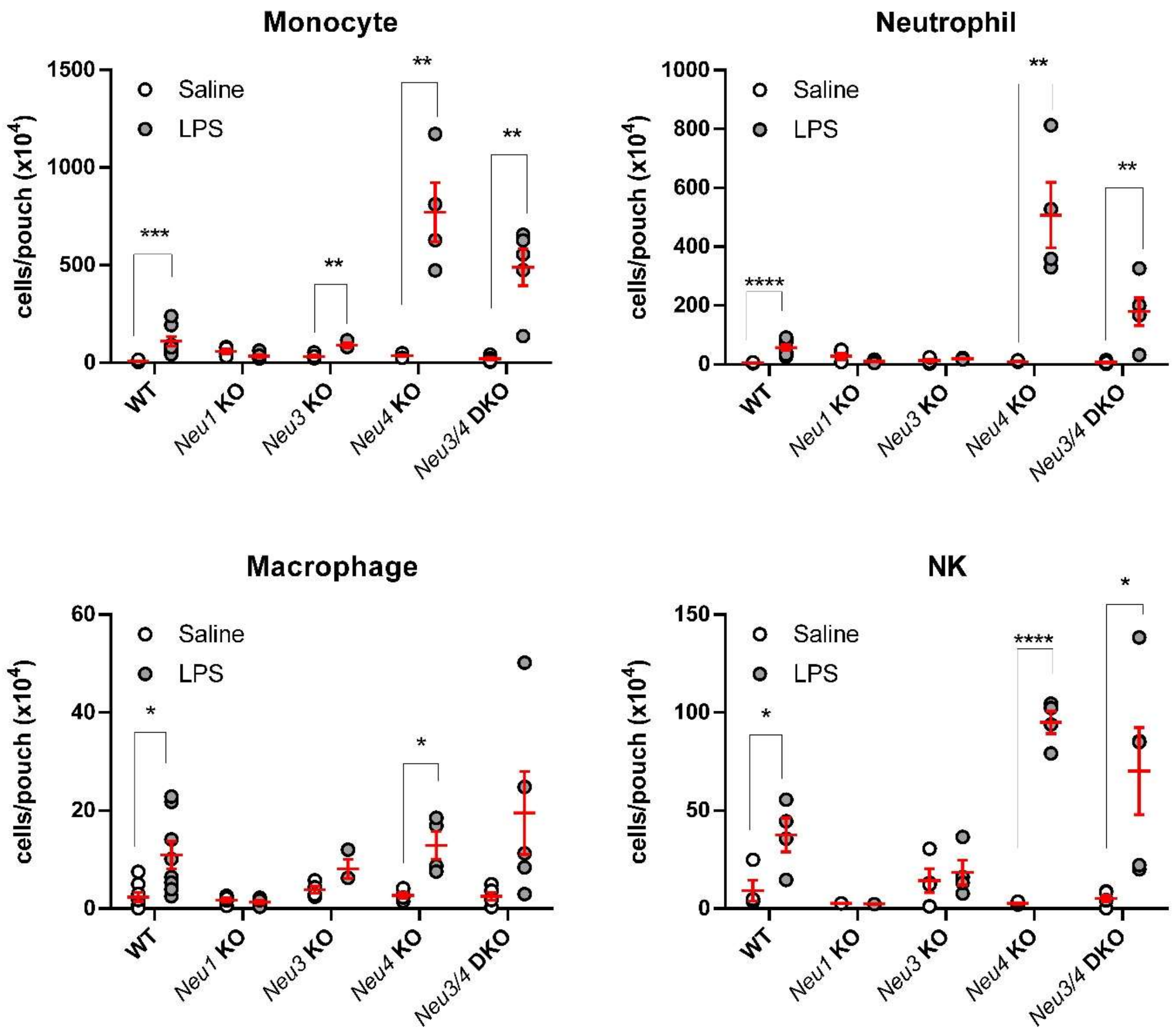

**Figure SI1: Raw cell counts of leukocyte subsets in the air pouch model.** Leukocytes collected from animals after saline or LPS treatment were counted by FACS and identified based on antibody markers. Cell types shown are Monocytes, Neutrophils, Macrophages, and NK cells. WT, NEU1 KO, NEU3 KO, NEU4 KO and NEU3/4 DKO mice were used in this study. The air pouch exudate was collected after 9 h. The data is presented as mean ± SEM, and conditions were compared to control using two-way ANOVA followed by Bonferroni multiple comparison test (*, $p \leq 0.05$; **, $p \leq 0.01$; ***, $p \leq 0.005$; ****, $p \leq 0.0001$).

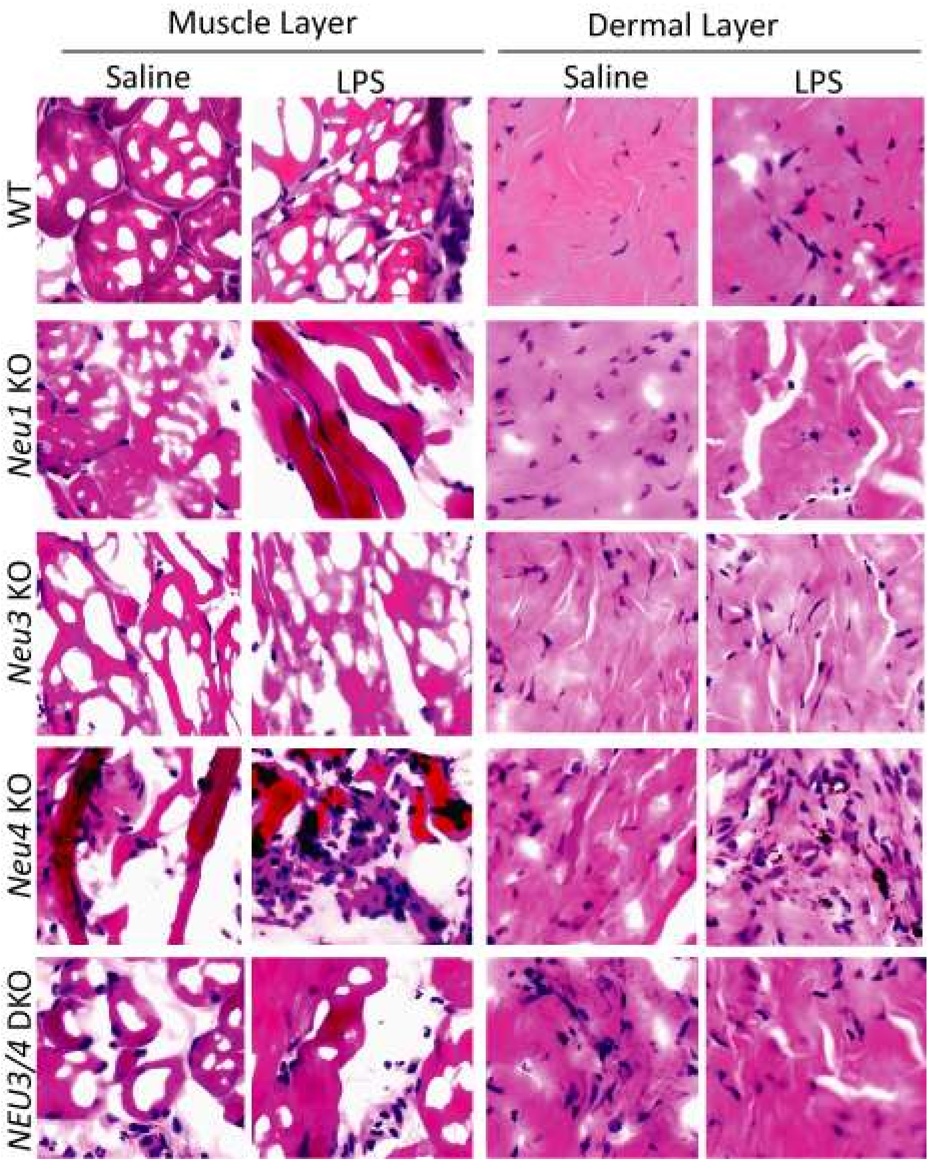

**Figure SI2: Immunohistochemistry of skin slices of the air pouch model.** Tissue from the air pouch model was collected, sectioned, and stained (H&E). Regions of tissue were identified as dermis or muscle. Representative images of the dermis and muscle regions of different mice groups. Random fields from each region were used to determine leukocyte counts shown in **Figure 3.**

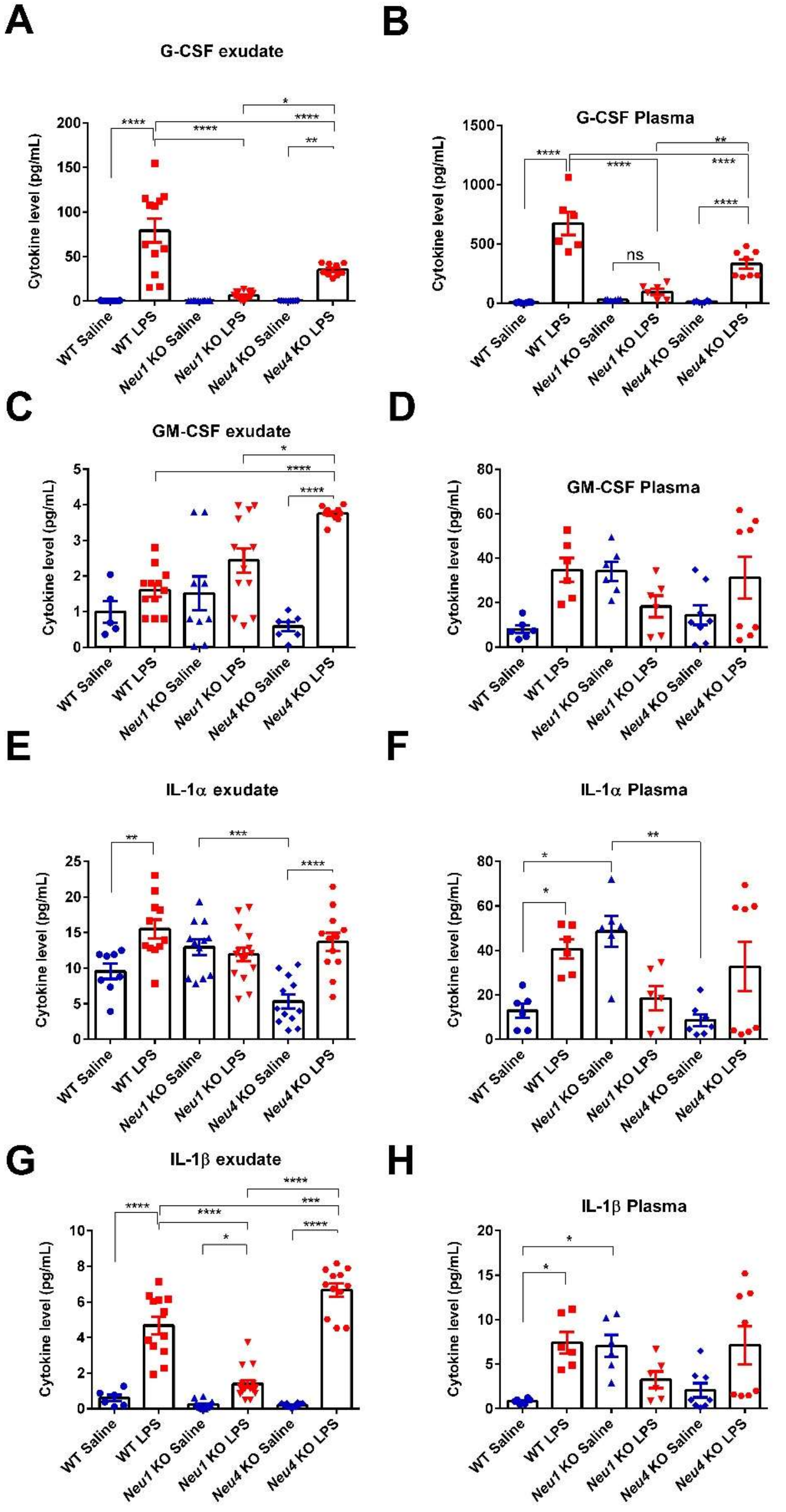
A
G-CSF exudate
B
G-CSF Plasma
ns
C
GM-CSF exudate
D
GM-CSF Plasma
E
IL-1α exudate
F
IL-1α Plasma
G
IL-1β exudate
H
IL-1β Plasma
Cytokine level (pg/mL)
WT Saline
WT LPS
Neu1 KO Saline
Neu1 KO LPS
Neu4 KO Saline
Neu4 KO LPS

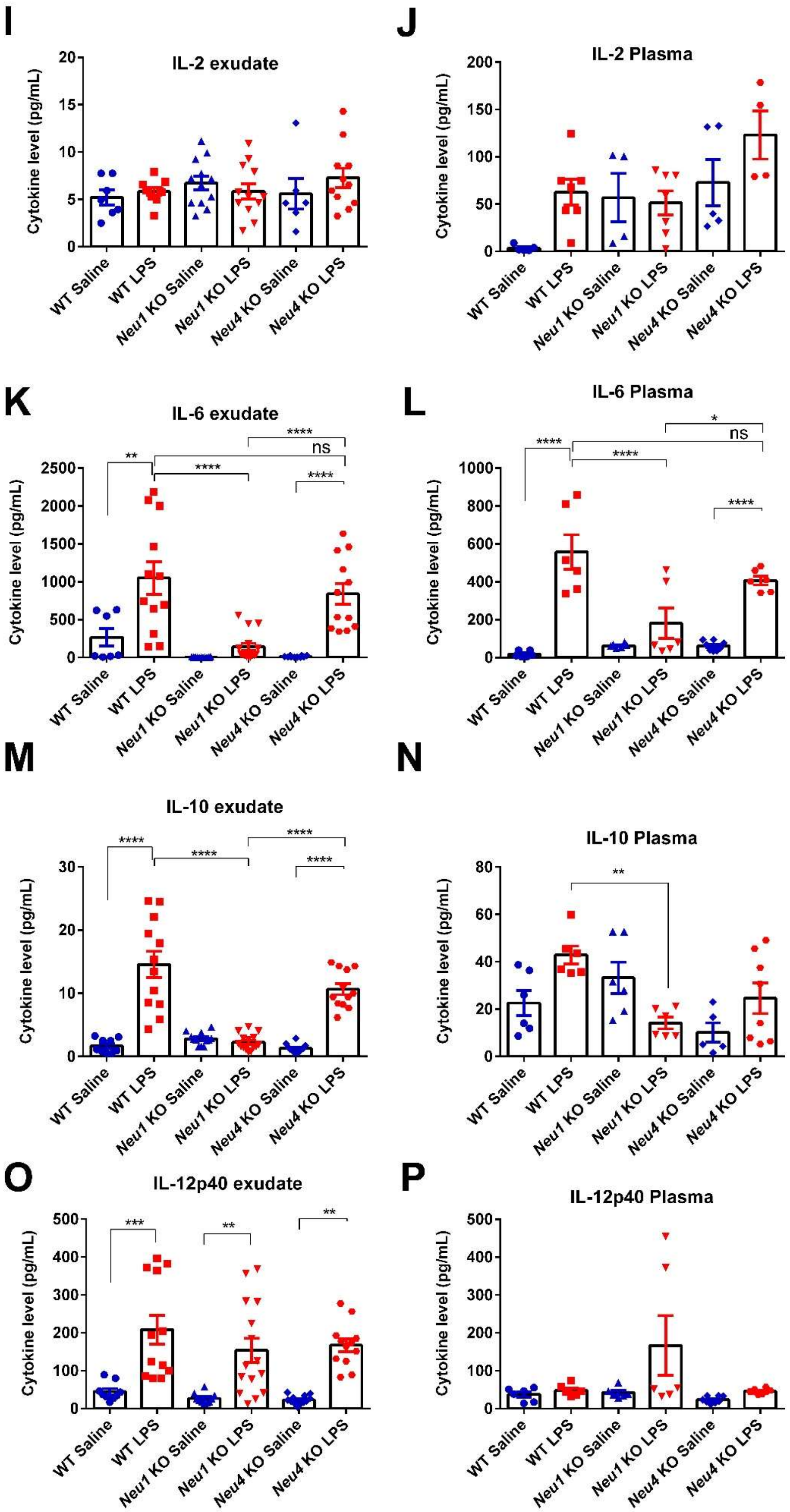
I
IL-2 exudate
J
IL-2 Plasma
K
IL-6 exudate
L
IL-6 Plasma
M
IL-10 exudate
N
IL-10 Plasma
O
IL-12p40 exudate
P
IL-12p40 Plasma
Cytokine level (pg/mL)
WT Saline
WT LPS
Neu1 KO Saline
Neu1 KO LPS
Neu4 KO Saline
Neu4 KO LPS
ns

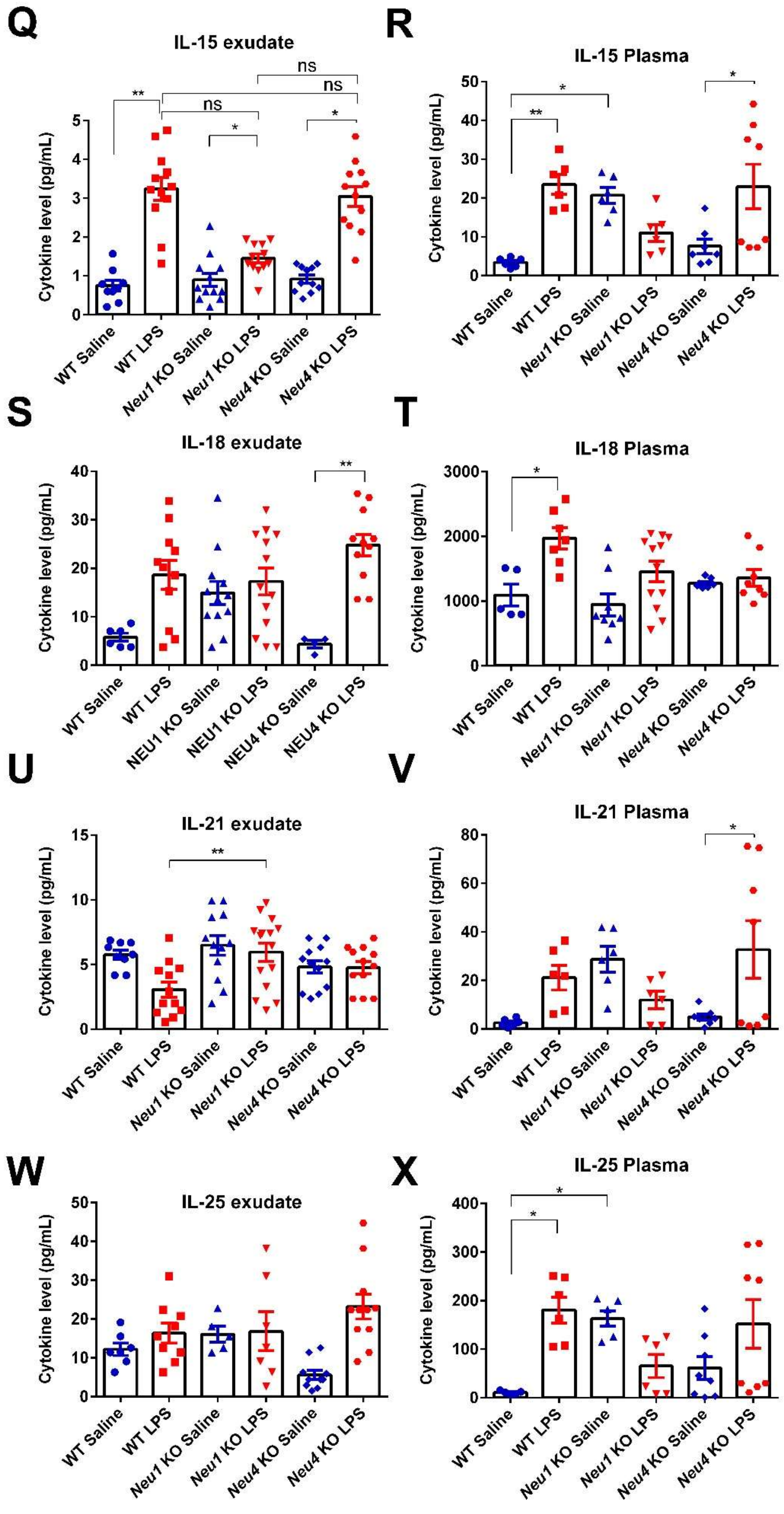
Q
IL-15 exudate
R
IL-15 Plasma
S
IL-18 exudate
T
IL-18 Plasma
U
IL-21 exudate
V
IL-21 Plasma
W
IL-25 exudate
X
IL-25 Plasma
Cytokine level (pg/mL)
ns
*
**
WT Saline
WT LPS
Neu1 KO Saline
Neu1 KO LPS
Neu4 KO Saline
Neu4 KO LPS
NEU1 KO Saline
NEU1 KO LPS
NEU4 KO Saline
NEU4 KO LPS

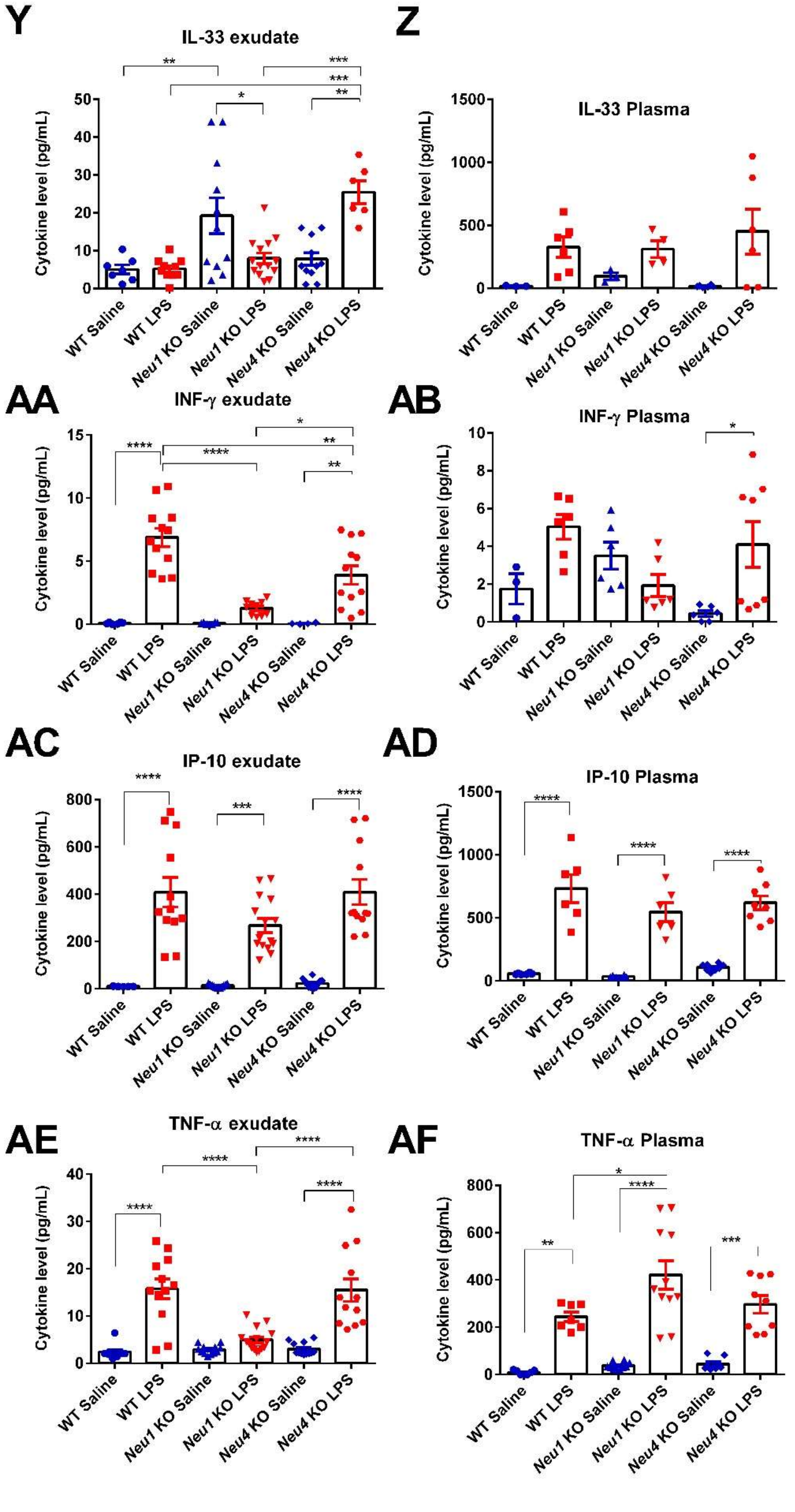
Y
IL-33 exudate
Z
IL-33 Plasma
AA
INF-γ exudate
AB
INF-γ Plasma
AC
IP-10 exudate
AD
IP-10 Plasma
AE
TNF-α exudate
AF
TNF-α Plasma
Cytokine level (pg/mL)
WT Saline
WT LPS
Neu1 KO Saline
Neu1 KO LPS
Neu4 KO Saline
Neu4 KO LPS

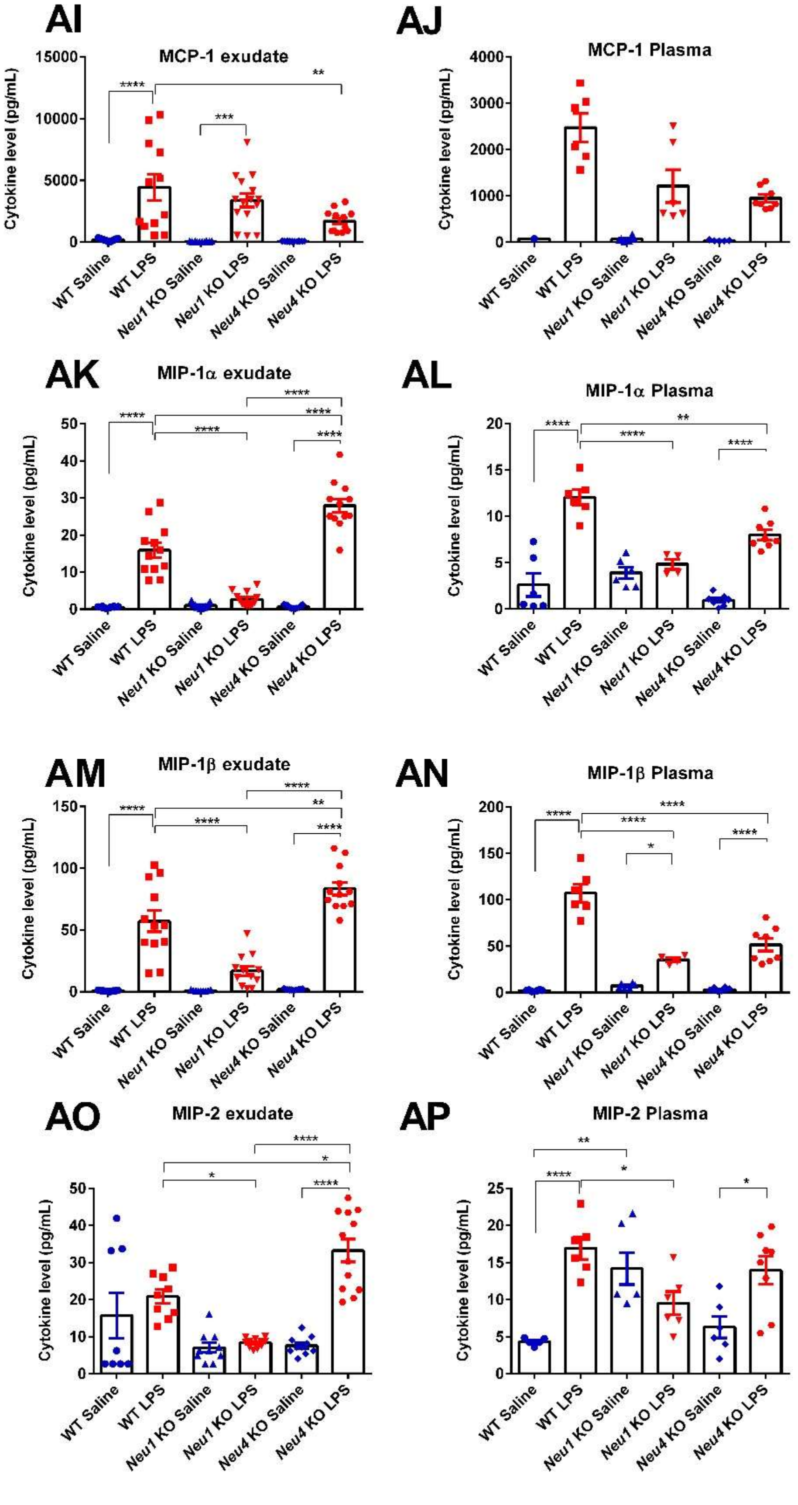
AI
MCP-1 exudate
Cytokine level (pg/mL)
AJ
MCP-1 Plasma
AK
MIP-1α exudate
AL
MIP-1α Plasma
AM
MIP-1β exudate
AN
MIP-1β Plasma
AO
MIP-2 exudate
AP
MIP-2 Plasma
WT Saline
WT LPS
Neu1 KO Saline
Neu1 KO LPS
Neu4 KO Saline
Neu4 KO LPS

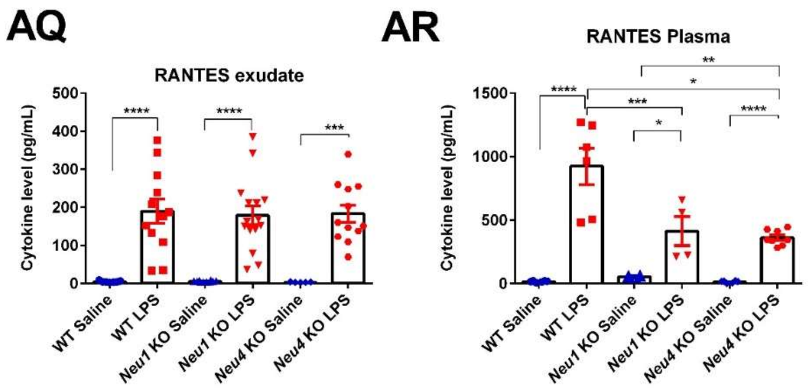

**Figure SI3. Cytokine levels in mice treated with LPS.** Cytokines were analyzed using the Autoplex Analyser CS1000 system (Perkin Elmer, Waltham, MA) with a commercial ProcartaPlex Mouse Cytokine Panel Assay kit (Thermo Fisher Scientific Inc., Rockford, USA) in accordance with the manufacturer's instruction. Levels for cytokines from exudate and plasma samples are shown for: (**A**-**B**) G-CSF, (**C**-**D**) GM-CSF, (**E-F**) IL-1α, (**G-H**) IL-1β, (**I-J**) Il-2, (**K-L**) IL-6, (**M-N**) IL-10, (**O-P**) IL-12p40, (**Q-R**) IL-15, (**S-T**) IL-18, (**U-V**) IL-21, (**W-X**) IL-25, (**Y-Z**) IL-33, (**AA-AB**) INF-γ, (**AC-AD**) IP-10, (**AE-AF**) TNF-α, (**AI-AJ**) MCP-1, (**AK-AL**) MIP-1α, (**AM-AN**) MIP-1β, (**AO-AP**) MIP-2, and **(AQ-AR**) RANTES. Bars represent the mean values ± standard error of the mean (SEM) in units of pg/mL. Samples were compared to controls using a one-way ANOVA followed by Tukey's multiple comparison test (*, $p < 0.05$; **, $p < 0.01$; ***, $p < 0.001$; ****, $p < 0.0001$).

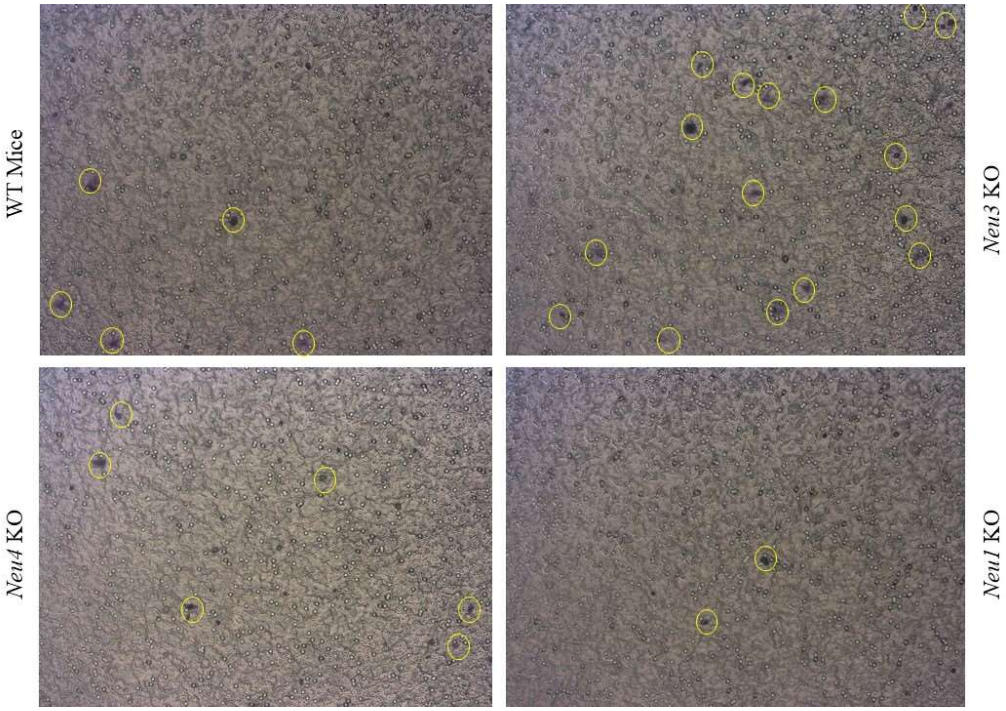

**Figure SI4: Representative images from transmigration experiments.** Macrophages were isolated and differentiated from bone marrow of WT, NEU1 KO, NEU3 KO, and NEU4 KO mice. Cells (2.5 x$10^4$) were placed in the upper chamber of the transmigration plate and the experiment was carried out for 5 h. The number of cells that infiltrated the FN-coated membrane was determined by counting of stained cells. The number of cells that infiltrated the FN-coated membrane was determined by counting of stained cells. A representative set of brightfield images is shown in the upper panel using a 10 X objective. Cell counts are shown in **Figure 6.**

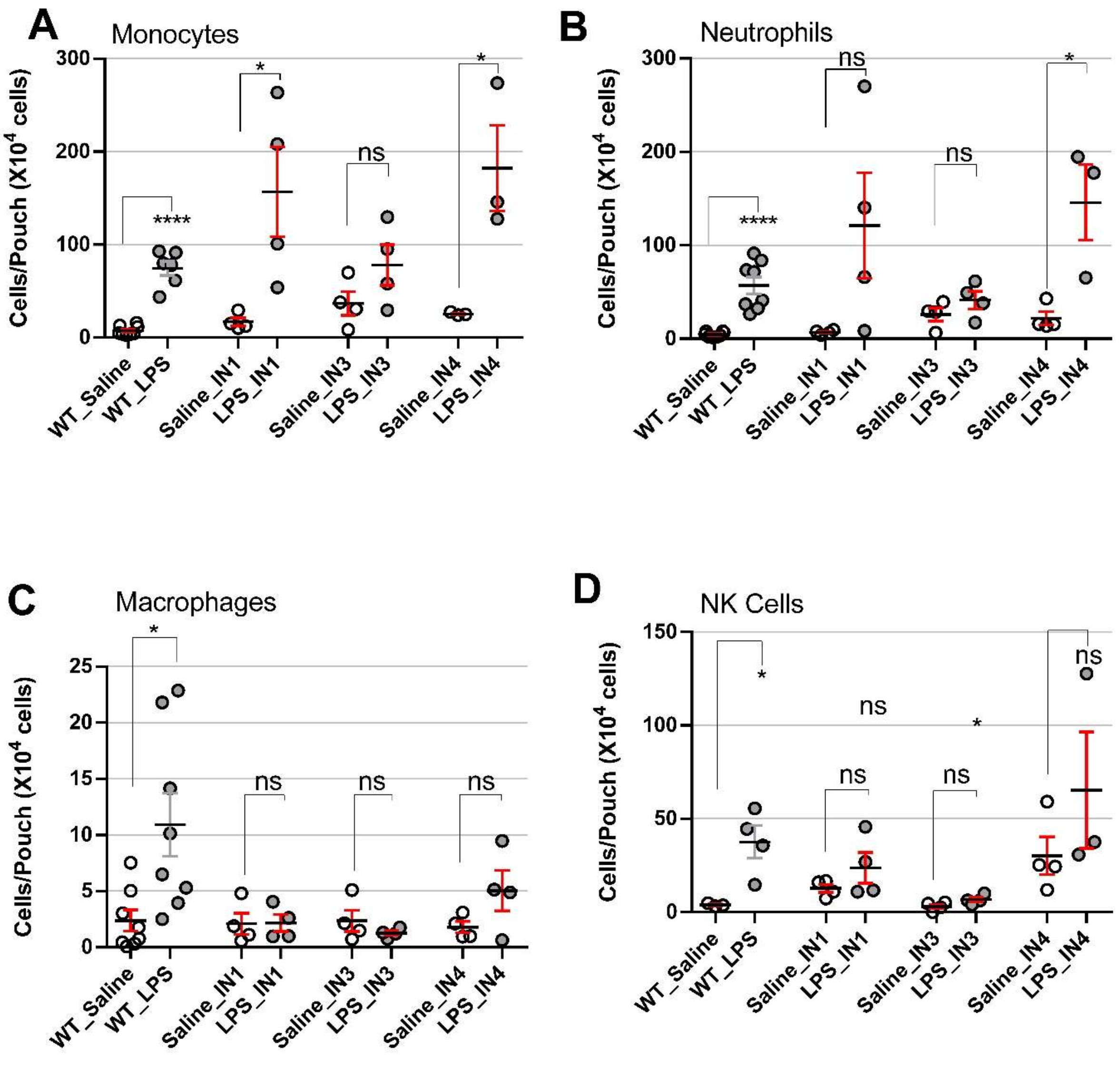

**Figure SI5. Leukocyte subsets in the air pouch model after treatment with neuraminidase inhibitors.** Leukocytes collected from animals were counted by FACS and identified based on antibody markers (see Methods). Inhibitors that target NEU1 (**IN1**, CG14600), NEU3 (**IN3**, CG22600), and NEU4 (**IN4**, CY16600) were used. Cell types were A. Monocytes, B. Neutrophils, C. Macrophages, and D. NK cells. Animals were WT only and inhibitor treatments were at 1 mg/kg. The air pouch exudate was collected after 9 h. The data is presented as mean ± SEM, and conditions were compared to control using Student's t-test (*, $p \leq 0.05$; **, $p \leq 0.01$; ***, $p \leq 0.005$; ****, $p \leq 0.0001$).

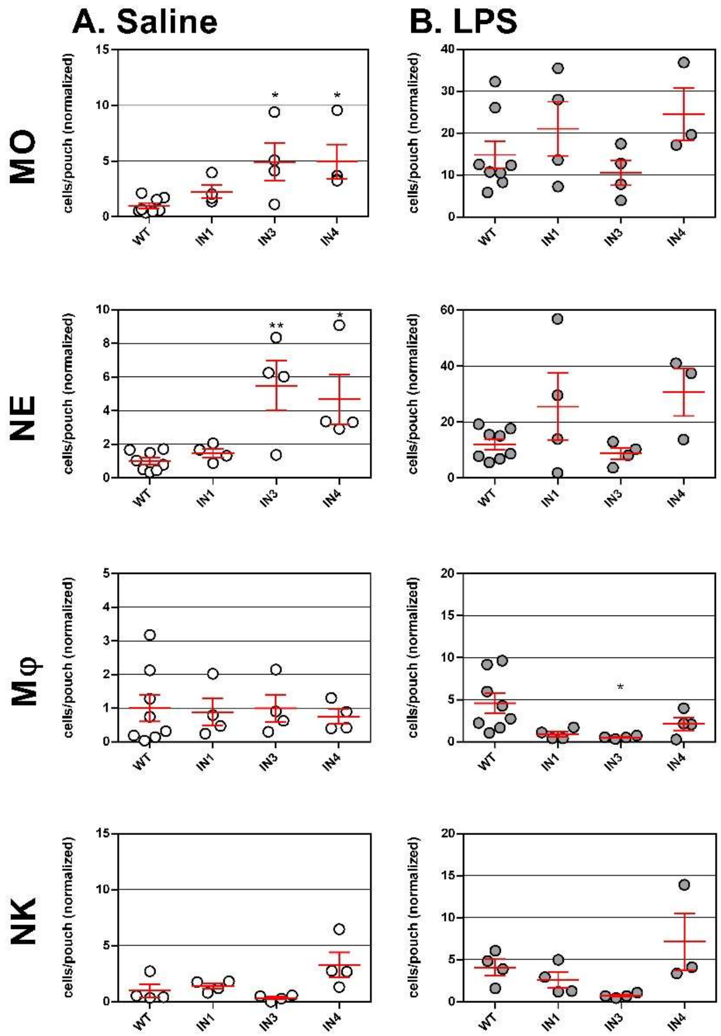

**Figure SI6. Leukocyte populations in the air pouch model after inhibitor treatment.** Leukocytes from mice treated with either treated with saline or neuramindase inhibitors (**IN1**, **IN3**, and **IN4**). After which animals were either injected with saline (○), or LPS (●) treatment. Collected leukocytes were processed with marker antibodies and were counted by flow cytometry. The cells were identified after staining with marker-specific fluorochrome-conjugated antibodies as monocyte (MO), neutrophil (NE), macrophage (MΘ), or natural killer (NK) cells. The air pouch exudate was collected after 9 h. The data is presented as cell counts (Mean ± SEM) compared to the respective WT saline controls. Means were compared to control using one-way ANOVA following Dunnet's t-test (*, $p \leq 0.05$; **, $p \leq 0.01$; ***, $p \leq 0.005$; ****, $p \leq 0.0001$). Raw cell counts are presented in Fig SI5.
